# Bioactive coatings on 3D printed polycaprolactone scaffolds for bone regeneration: a novel murine femur defect model for examination of the biomaterial capacity for repair

**DOI:** 10.1101/2023.12.15.569064

**Authors:** Karen. M. Marshall, Jonathan P. Wojciechowski, Vineetha Jayawarna, Abshar Hasan, Cécile Echalier, Sebastien J. P. Callens, Tao Yang, Janos M. Kanczler, Jonathan I. Dawson, Alvaro Mata, Manuel Salmeron-Sanchez, Molly M. Stevens, Richard O.C. Oreffo

## Abstract

Bone tissue engineering is a rapidly advancing field that seeks to develop efficacious approaches for treating non-healing fractures and large bone defects. Healing complications arise due to trauma, disease, infection, aseptic loosening of orthopaedic implants or iatrogenic causes. An ideal biodegradable scaffold would induce and support bone formation until the bone matrix is sufficiently stable to facilitate healing. The current study has examined bone augmentation, using functionalised coated scaffolds, with the hypothesised potential to induce skeletal cell differentiation for the repair of critical-sized bone defects. However, challenges in clinical translation arise from the alterations in cellular microenvironment that are present in the translation from *in vitro* to *in vivo* with the application of animal models of progressively increasing size and complexity of the implantation site. 3D printed, porous poly(caprolactone) trimethacrylate (denoted PCL-TMA900) scaffolds were applied within a murine femur defect, stabilised by a polyimide intramedullary pin, to assess the efficacy of select coatings in inducing bone formation. The PCL-TMA900 scaffolds were coated with i) elastin-like polypeptide (ELP), ii) poly(ethyl acrylate)/fibronectin/bone morphogenetic protein-2 (PEA/FN/BMP-2), iii) both ELP and PEA/FN/BMP-2 concurrently, or iv) Laponite™ nanoclay binding BMP-2, as bioactive coatings. The murine femur defect model was refined to assess the coated PCL-TMA900 scaffolds in an osseous defect, with sequential microcomputed tomography (µCT) and histological analysis of the new bone tissue.

Overall, PCL-TMA900 was found to be an optimal robust, biocompatible, 3D printable scaffold material. All PCL-TMA900 scaffolds, uncoated and coated, showed integration with the femur. The PCL-TMA900 scaffold coated with the nanoclay material Laponite™ and BMP-2 induced consistent, significant bone formation compared to the uncoated PCL-TMA900 scaffold. Bone formation was observed within the pores of the Laponite/BMP-2 coated scaffold. Critically, no heterotopic bone formation was observed as the BMP-2 was retained around the scaffold and not released into the tissues, producing bone around the scaffold in the desired shape and volume, compared to bone formation observed with the positive control (collagen sponge/BMP-2 construct). In comparison, the ELP coated and PEA/FN/BMP-2 scaffolds did not demonstrate significant or consistent bone formation compared to uncoated PCL-TMA900 control scaffolds.

In summary, nanoclay Laponite™/BMP-2 coated PCL-TMA900 scaffolds offer a biodegradable, osteogenic construct for bone augmentation with potential for development into a large scale polymer scaffold for translation to the clinic.

## Introduction

Bone tissue engineering has evolved due to the critical need for innovative constructs to replace grafted bone or suboptimal materials currently applied in clinical practice. Defects arising in bone may occur due to trauma, fracture non-union, infection, resection in oncology cases or congenital conditions [1]. In certain clinical cases, amputation is required as the limb is unsalvageable due to significant soft tissue, vascular and/or nerve damage [2]. Bone is the second most grafted tissue after blood, frequently using the patient’s own bone (autograft) from the iliac crest of the pelvis [3, 4]. From a clinical perspective, autograft is the optimal material given it is non-immunogenic, facilitates osteoinduction due to the presence of growth factors, osteoconduction as bone is a 3D scaffold, and osteogenesis due to the cellular component within [5]. However, there are a number of significant limitations including donor site complications such as infection, pain and, naturally, limited autograft volume [6]. Thus, the ideal bone substitute material is biocompatible, bioresorbable, osteoconductive, osteoinductive, porous with a comparable structure and strength to bone, easy to use and cost effective [1]. Critically, the material must be able to withstand the complex load and mechanical forces applied, such as weight bearing or mastication [7].

We have previously assessed the biocompatibility of a poly(caprolactone) trimethacrylate (PCL-TMA900) scaffold material functionalised with a variety of scaffold coatings including elastin-like polypeptides (ELPs), poly(ethyl acrylate)/fibronectin/bone morphogenetic protein-2 (PEA/FN/BMP-2) and the nanoclay Laponite™, *in vitro*, using the chorioallantoic membrane (CAM) assay and in a murine subcutaneous implantation model [8, 9]. ELPs are recombinantly expressed polypeptides based on the properties of elastin and are highly biocompatible as these polypeptides are non-immunogenic [10]. The ELP matrix has been previously reported to sequester calcium and phosphate ions from the surrounding environment to nucleate apatite nanocrystals [11]. ELP matrix can also be applied as a coating on hard scaffolds to facilitate bone formation and osseointegration *in vivo* [12]. The poly(ethyl acrylate) (PEA), fibronectin (FN) and bone morphogenetic protein-2 (BMP-2) coating was developed to permit PEA to induce a fibrillar arrangement of FN, exposing FN binding sites to present BMP-2 to adjacent skeletal cells, enhancing osteogenic differentiation while utilising low concentrations of BMP-2 [13]. The nanoclay, Laponite™, has been shown to sequester growth factors, such as vascular endothelial growth factor or BMP-2, allowing targeted delivery to the requisite site with a subsequent angiogenic or osteogenic tissue response induced [8, 14–16].

In our previous study, only the Laponite coating, when used to bind BMP-2, was effective at inducing robust ectopic bone formation and osteogenic differentiation *in vitro* [8]. Previous work however, indicated that the ELP and PEA/FN/BMP-2 coatings may function optimally in the osteogenic environment of an orthotopic site – such as in a model of fracture repair [9, 12, 13, 17]. Therefore, application of all the coatings in a fracture model was performed to determine the efficacy of each coating in promoting bone repair. In these studies, we use an optimised preclinical critical-sized mouse femur defect model undertaking sequential data collection over the 8 week study period.

Critical-sized femur defects have been used in rodents to assess bone formation in a plethora of studies [18]. Thus, in mice, plates with screws, external skeletal fixators (ESFs), and variations in defect length from 2-4 mm depending on the type of mouse have been examined [19–23]. In studies of the fractured bone, fracture gaps of 0.1 mm to 2 mm were maintained by an ESF in the mouse femur, however, a defect of 3 mm is deemed necessary to be of critical size [24, 25]. A femoral metaphyseal defect model in mice has been described in the examination of bone healing, although in the current work, such a defect model would have been too small to accept a solid 3D printed, porous scaffold [26]. Intramedullary (IM) pinning was the chosen method of stabilisation of the scaffolds within the femur defect [27]. This was a refined method as the pin did not exit the femur proximally and distally, limiting injury to the mice and damage to the medullary canal, with the use of a shorter pin centrally within the scaffold, retaining the scaffold in place (**Supplementary Figure 1**).

In an initial pilot study, poly(caprolactone) scaffolds and collagen sponge loaded with BMP-2 as a positive control stabilised using a metal pin were trialled. The hypothesis was that the scaffolds would be of sufficient length (5 mm) that non-union of the fracture gap would result while the collagen sponge containing BMP-2 would unite the femur ends. These key points were established in a pilot study (**Supplementary Figures 2 and 3)**. In a second pilot femur defect study, the use of PCL-TMA900 scaffolds, stabilised with a hypodermic needle (25G) or polyimide/plastic tubing in the medullary canal allowed assessment of the biocompatibility of the PCL-TMA900. In addition, the welfare, imaging and histological analysis benefits and potential complications of using a plastic pin in comparison to a metal pin were examined (**Supplementary Figure 4**). The radiolucent plastic pin was found to be suitable for IM fixation, microcomputed tomography (µCT) imaging and histological analysis, with, notably, negligible destruction of the animal tissues as pin removal was not required.

Thus, the current study has examined bone formation on coated PCL-TMA900 scaffolds in comparison to the uncoated PCL-TMA900 scaffold (negative control) and BMP-2 soaked collagen sponge scaffold (positive control). The hypothesis under examination was that at least one of the coated (ELP, PEA/FN/BMP-2, ELP/PEA/FN/BMP-2, or Laponite/BMP-2) scaffolds would induce significantly greater bone formation in the defect site over an 8 week period compared to the uncoated control, as assessed by µCT imaging and histology.

## Materials and Methods

### Materials

Reagents were purchased as follows: Ethyl acrylate (E9706, Sigma, UK), ELP with statherin sequence (SN_A_15) (Technical Proteins Nanobiotechnology, Valladolid, Spain); recombinant human BMP-2 (Infuse/InductOS Bone graft kit, Medtronic, USA); recombinant human fibronectin (R&D systems, Biotechne, UK); Laponite (BYK); phosphate buffered saline (PBS) (Scientific Laboratory Supplies, SLS); Alcian blue 8X, Light green SF, Orange G 85% pure, paraformaldehyde 96% extra pure, phosphomolybdic acid hydrate 80% (Acros Organics); Picrosirius Red, Van Gieson’s stain, Weigert’s Haematoxylin Parts 1 and 2 (Clintech Ltd, UK); acetone, acid fushsin, histowax, ponceau xylidine, silver nitrate, sodium hydroxide pellets, sucrose, glycine, L-glutamic acid, sodium chloride, polysorbate 80 (Merck, UK); dibutyl phthalate xylene (DPX), Histoclear (Thermofisher Scientific, UK); Fast green and sodium thiosulphate (VWR); Lubrithal (Dechra, UK), Isoflurane (Dechra, UK), Buprenorphine (Buprecare® multidose, Animalcare, UK), and Vetasept® sourced from MWI animal health, UK. 5/0 PDS II suture (Ethicon, USA) from NHS supply chain. All other consumables and reagents were purchased from Sigma-Aldrich, UK.

### Mouse type and housing

All procedures were performed in accordance with institutional guidelines, with ethical approval and under PPL P96B16FBD and PPL PP8873619, in accordance with the regulations in the Animals (Scientific Procedures) Act 1986 and using the ARRIVE guidelines. Forty-two, adult, male, MF-1, wild type mice, bred on site and group housed in individually ventilated cages (IVCs) were used. Mice had access to *ad libitum* standard pellets and water. All mice were healthy with similar mean weight per group (mean and standard deviations to 1 decimal place of weight for each group were: uncoated scaffold 49.9 ± 4.1 g, ELP coated scaffold 49.9 ± 6.3 g, PEA/FN/BMP-2 coated scaffold 49.6 ± 5.1 g, ELP/PEA/FN/BMP-2 coated scaffold 50.2 ± 8.7 g, Laponite coated scaffold 52.2 ± 3.6 g, Laponite/BMP-2 coated scaffold 49.2 ± 7.4 g, collagen sponge/BMP-2 construct 51.0 ± 7.1 g), with normal mobility prior to surgery.

### Synthesis of the PCL-TMA900 polymer

PCL-TMA of this molecular weight has been synthesised and 3D printed using mSLA as previously detailed [28–30]. The PCL-TMA900 polymer was synthesised and characterised as detailed in Marshall *et al.* [8] using poly(caprolactone) triol, M_n_ = 830 Da as a starting material (Boc Sciences). Poly(caprolactone) triol (100 g, 0.12 mmol, 1 eq), anhydrous dichloromethane (300 mL) and triethylamine (100 mL, 0.72 mmol, 6 eq) were added to a 1 L two-necked round bottom flask. The reaction was placed under a nitrogen atmosphere and then cooled in an ice-water bath for 15 minutes. A pressure-equalising dropper funnel charged with methacryloyl chloride (53 mL, 0.67 mmol, 4 eq) was attached to the round bottom flask. The methacryloyl chloride was added dropwise over approximately 3 hours. The reaction was covered with aluminium foil to protect it from light and allowed to stir and warm to room temperature (RT) overnight. The next day, methanol (50 mL) was added to quench the reaction, which was allowed to stir at RT for 30 minutes. The reaction mixture was dissolved in dichloromethane (800 mL), transferred to a separating funnel and washed with 1 M aqueous hydrochloric acid solution (5×250 mL), saturated sodium bicarbonate solution (4×400 mL) and brine (1×400 mL). The organic layer was then dried with anhydrous magnesium sulphate, filtered and concentrated via rotary evaporation. The crude yellow liquid was then purified using a silica plug, with dichloromethane as the eluent. Fractions containing PCL-trimethacrylate were pooled and concentrated via rotary evaporation. The PCL-trimethacrylate was transferred to a brown glass vial and dried using a stream of air (through a plug of CaCl_2_) overnight to yield the title compound as a slightly yellow viscous liquid (94.25 g). The PCL-trimethacrylate was supplemented with 200 ppm (w/w) of 4-methoxyphenol (MEHQ) as an inhibitor.

^1^H NMR (400 MHz, CDCl_3_) δ 6.11 – 6.05 (m, 3H), 5.61 – 5.50 (m, 3H), 4.16 – 3.99 (m, 18H), 2.34-2.27 (m, 12H), 1.96 – 1.88 (m, 9H), 1.71 – 1.60 (m, 26H), 1.47 – 1.31 (m, 12H), 0.98 – 0.83 (m, 3H).

The characterisation data agrees well with that previously reported [28]. The degree of functionalisation was determined as >90% [8].

### Scaffold design and 3D printing of the PCL-TMA900 scaffolds

The poly(caprolactone) trimethacrylate (PCL-TMA900) scaffolds were designed using a custom Matlab script, based on the Schwarz P triply periodic minimal surface [31]. This resulted in a bicontinuous, porous, tubular shape (3 mm diameter, 5 mm height, pore size 0.5 mm and surface area 113.9 mm^2^), with a central hole of 0.6 mm diameter for the IM pin (**Supplementary Figure 5**).

The PCL-TMA900 scaffolds were printed using masked SLA 3D printing on a Prusa SL1. The resin was prepared for 3D printing by initially dissolving 0.1% (w/w) 2,5-thiophenediylbis(5-tert-butyl-1,3-benzoxazole) (OB+) as a photoabsorber in the PCL trimethacrylate by stirring at RT for 1 hour. Finally, 1.0% (w/w) diphenyl(2,4,6-trimethylbenzoyl)phosphine oxide (TPO-L) as a photoinitiator was added to the resin. The scaffolds were sliced in PrusaSlicer with their longitudinal axis parallel to the surface of the build platform. Supports and pads were used to raise the objects off the build platform, with ten supports placed along the scaffold to ensure adhesion to the build platform during the printing whilst not compromising the structure. The scaffolds were sliced at a layer height of 50 µm using a 25 second exposure time per layer.

After printing, the scaffolds were rinsed with ethanol and removed from the build plate. The support structures were removed from the scaffolds with a fine sprue cutter, then the scaffolds were sonicated in ethanol (5 × 5 minutes) and allowed to dry for 15 minutes at RT. The scaffolds were post-cured using a Formlabs Form Cure for 60 minutes at RT. After post-curing, the central hole of the scaffolds was cleared by hand with a 0.6 mm drill bit and rotation motion to ensure good fit with the pins. The scaffolds were then soaked in ethanol overnight at RT on a rocker (100 rpm), washed with ethanol (3×) and allowed to dry at RT before being ELP and/or PEA coated then sterilised by ethylene oxide (EO), or left uncoated and EO sterilised as the uncoated scaffold control group.

### Intramedullary Pin preparation

EO sterile, 1.2 cm long polyimide tubing (internal diameter 0.0136″, outer diameter 0.0231″, wall thickness 0.00475″, Microlumen, USA) was used as the IM pin in every mouse (**Supplementary Figure 1)**. Initially, a metal pin was applied in the first pilot study, however, the metal pin produced µCT imaging artefacts, stress shielding and bone resorption (**Supplementary Figures 2 and 3**). The polyimide/plastic pin was verified as suitable for use by implantation into cadaver mice and findings in the second pilot study which compared the use of a metal in contrast to a polyimide pin (**Supplementary Figure 4**). The plastic pin was found to be adequate and allowed bone quantification without µCT imaging artefacts.

### Preparation of the uncoated and coated PCL-TMA900 scaffolds

The following scaffolds were prepared: a negative control group of uncoated PCL-TMA900 scaffolds (n=6), together with ELP, PEA/FN/BMP-2, ELP/PEA/FN/BMP-2, Laponite alone or Laponite/BMP-2 coated PCL-TMA900 scaffolds (n=7 scaffolds prepared for each group including a spare scaffold).

The scaffolds were sterilised by EO prior to use as uncoated scaffolds, prior to the application of Laponite or after PEA or ELP coating. Prior to use, a 0.6 mm drill bit (KATUR, Amazon website) was autoclaved at 126 °C and inserted by hand, with rotation, into the central hole of each scaffold to ensure easy insertion of the polyimide tubing. In a class 2 hood, the scaffolds were threaded onto the polyimide tubing by hand wearing sterile nitrile gloves and placed in a 6 well plate to maintain sterility.

#### PEA coating application

PEA coating of the scaffolds was performed using a custom-built plasma chamber as previously described [32]. In brief, the scaffolds were treated in air plasma for 5 minutes before being exposed to monomer plasma. Plasma polymerization of the ethyl acrylate monomer was carried out in a custom-made capacitively coupled plasma installation for low-pressure plasma in a 15-L T-shaped reactor made of borosilicate and stainless-steel end plates sealed with Viton O-rings. A vacuum was produced by a rotary pump or a scroll pump (both BOC Edwards, UK), with operating experiment pressures for the monomer plasma from 0.15 to 0.25 mbar. The plasma was initiated using two capacitively coupled copper band ring electrodes situated outside of the reactor chamber and connected to a radio frequency power supply (Coaxial Power System Ltd.) at 50 W for 15 minutes. The monomer pressure was controlled using speedivalves (BOC Edwards, UK) and monitored with a pirani gauge (Kurt J. Lesker).

#### ELP coating application

ELP coating of scaffolds was based on previous published methods [12]. In brief, lyophilized ELP powder was dissolved in a solvent mixture of DMF/DMSO (at 9/1 ratio, v/v) to prepare a 5% (w/v) ELP solution followed by addition of hexamethyl diisocyanate (HDI) crosslinker (cross-linker to lysine ratios of 12/1). 3D printed PCL-TMA900 scaffolds were immersed in the ELP solution for 10–15 seconds and left to dry overnight at RT (22 °C) within a glovebox (BELLE Technology, UK) maintained at a humidity <20%. Dry ELP coated scaffolds were subsequently washed several times with deionised water (dH_2_O) to remove excess HDI and termed as ELP coated scaffolds [12]. The ELP coating was confirmed to be present on the PCL-TMA900 scaffold surface using scanning electron microscopy [12]. For the ELP/PEA/FN/BMP-2 coated scaffolds, the ELP coating was applied as described above, followed by PEA. Finally, the FN and BMP-2 were adhered to the scaffolds as described below.

#### Application of FN and BMP-2 to the PEA coating

The method of coating has been described in Marshall *et al.* [9]. In brief, on the day of surgery, the PEA and ELP/PEA coated scaffolds were immersed in FN (20 µg/mL) followed by InductOs® BMP-2 (5 µg/mL) with 1 scaffold per vacutainer and 1 mL of solution used per vacutainer using PBS (with CaCl_2_ and MgCl_2_) as the diluent, under vacuum for 1 hour at RT. The scaffolds were rinsed in PBS with CaCl_2_ and MgCl_2_ and moved to a new vacutainer between FN and BMP-2 adsorption. The scaffolds were given a final rinse in PBS (with CaCl_2_ and MgCl_2_) before threading onto the polyimide tubing.

#### Laponite and BMP-2 coating

The method of coating has been described in Marshall *et al.* [8]. In brief, a 1 % (wt/wt) Laponite solution was prepared and stored at 4°C, 48 hours prior to use. 0.2 g Laponite was dispersed, under stirring in 19.8 g dH_2_O and the resulting clear suspension autoclaved to sterilise. After autoclaving, the weight of lost dH_2_O was replaced with sterile dH_2_O. On the day of surgery, 14 PCL-TMA900 scaffolds were immersed individually in 1 mL of 1% (w/w) Laponite solution. After 1 hour at RT in the sterile class II hood, the scaffolds were removed with sterile forceps and threaded onto 25-gauge needles in a 24-well plate (**Supplementary Figure 6**) facing upwards to ensure the scaffolds dried without risk of transfer of coating from the scaffold. The excess Laponite was gently removed with a 100 µL pipette tip. The scaffolds were allowed to dry for 2 hours prior to use as a Laponite only coated scaffold (n=7) or underwent further coating in BMP-2 (n=7).

For BMP-2 binding, seven Laponite coated PCL-TMA900 scaffolds were immersed individually in 1 mL of 5 µg/mL BMP-2 (InductOs® BMP-2) in PBS (not containing CaCl_2_ and MgCl_2_) in 1.5 mL low protein bind Eppendorf tubes at RT for 24 hours. After 24 hours, the PCL-TMA900 scaffolds were taken out of the BMP-2 solution and dried on needles in a category II hood. Polyimide tubing was threaded through the central hole of the scaffold prior to implantation.

#### Collagen Sponge/BMP-2 construct

Collagen sponges loaded with 5 µg of InductOs® BMP-2 was used as the positive control scaffold. The collagen sponge was EO sterile but to ensure sterility, the collagen sponge was UV sterilised overnight in a sterile class 2 hood. Approximately 20, 3 mm × 3 mm collagen sponge pieces were threaded onto a 23-gauge needle. The BMP-2 was diluted with InductOs® buffer (**Supplementary Table 1**) to 333.33 µg/mL concentration and 15 µL of solution (5 µg mass of BMP-2) was added to the collagen sponge. With fluid addition, the collagen was moulded into a single 5 mm length construct. The construct was then dried for 2 hours to form a rigid 5 mm long scaffold and was then transferred from the needle to the polyimide tubing.

### Surgical procedure

General anaesthesia was induced in mice with an induction chamber using Isoflurane (5%) and 100% oxygen (0.8 L/min flow rate). The mouse was moved to the nose cone for maintenance of a surgical plane of anaesthesia at approximately 2-2.5% isoflurane and 100% oxygen (0.8 L/min flow rate). Lubrithal (Dechra, UK) was applied to the eyes, ear marking was performed for identification, the entire left hindlimb and side to mid-body of the mouse was shaved, and chlorhexidine/alcohol solution applied. Buprenorphine (0.05 mg/kg) was administered by subcutaneous injection. In right lateral recumbency, the left hindlimb third trochanter landmark was palpated, and a 3 cm incision was made over the femur. The white band of fat and fascia over the femur was dissected to separate the vastus lateralis and biceps femoris muscles (**Supplementary Figure 8 A)**. Care was taken not to probe medially or caudally to avoid injury to the femoral artery and sciatic nerve respectively. The femur was isolated using a micro spatula placed underneath to protect the soft tissues and to act as a skin retractor (**Supplementary Figure 8 B and C)**. A rotating circular saw blade (botiss biomaterials, Germany) was used to make a partial cut at the proximal and distal extents of the osteotomy. The proximal extent was half-way along the third trochanter, with attempt to preserve as much gluteal muscle attachment as possible. The distal cut was at the distal extent of the third trochanter, where a white triangular mark in the bone tapers off, visible during surgery, whereby removing a 2-3 mm piece of bone (**Supplementary Figure 8 D)**. Irrigation with sterile saline during bone cutting prevented thermal necrosis and aided removal of bone debris. Bleeding was minimal from the musculature and marrow cavity. A 25-gauge needle with the bevel removed with wire stripping pliers, bent at 4 mm length and autoclaved, was used to clear 4 mm depth within the marrow cavity of the proximal and distal femur prior to insertion of the polyimide pin (**Supplementary Figure 7**). The polyimide pin within the scaffold/collagen sponge, was inserted into the proximal and distal portion of the femur (**Supplementary Figure 8 E and F**). The PCL-TMA900 scaffold or collagen sponge construct were assessed to be correctly positioned within the defect (**Supplementary Figure 8 G and Figure 9).** The muscle layer was closed with 2 simple interrupted sutures with 5/0 PDS (Ethicon, USA) (**Supplementary Figure 8 H**). The skin was closed in a simple interrupted, horizontal mattress suture pattern using 5/0 PDS (Ethicon, USA), with the knots towards the tail to prevent interference by the mouse (**Supplementary Figure 8 I**). Following µCT imaging, the mice were recovered in a warming cage (Thermacage, datesand, UK) post operatively until alert and mobile and returned to their IVC cage. Food was offered from the cage floor to prevent the necessity of standing on the back legs to reach the food hopper, and behaviour, mobility and nesting behaviour were monitored. Buprenorphine (0.05 mg/kg) was administered by subcutaneous injection 6-8 hours post-operatively and continued twice daily for 3 days. Of note, in the current study, the scaffold was firmly attached to the proximal femur with the pin held straight within the medullary canal and scaffold in this region, creating a pivot point for the plastic pin to bend and to be held there under the adductor muscle force, compared to the second pilot study where the plastic pin bent at the edge of the proximal femur and therefore remained straight within the scaffold and distal portion of the femur, thereby maintaining a relatively straight limb. Use of a barbed pin to grip the endosteum of the diaphysis could perhaps improve stability and placement of the intramedullary pin [33]. There was a change in direction of the greater trochanter of the femur, from a lateral position to a cranial position, in all mice in all studies performed, due to the unavoidable disruption of the surrounding musculature at surgery, however, this did not appear to interfere with locomotion.

### µCT procedure and analysis

Micro-CT was performed using a MILabs OI-CTUHXR preclinical imaging scanner (Utrecht, The Netherlands). General anaesthesia was induced in an induction chamber with 5% Isoflurane (VetTech) and 100% oxygen (0.8 L/min flow rate), followed by maintenance of general anaesthesia at approximately 1.5-2% isoflurane throughout imaging, with the oxygen rate constant at ∼1.0 L/min. BioVet software connected to the Milabs scanner permitted monitoring of respiratory activity (minimum of 60 breaths per minute to judge anaesthetic depth) and setting the temperature of the scanning bed to 34°C. Lubrithal (Dechra, UK) was applied to both eyes to prevent corneal desiccation. Mice were imaged from the mid back to the toes, typically 2 µCT bed positions were selected, with a total scan time of approximately 10 minutes per acquisition. The mice recovered on 100% oxygen, whilst being kept warm on the imaging bed. After signs of movement were observed, the mouse was transferred to a warming box (Thermacage, datesand, UK) prewarmed at 30°C until normal mouse activity resumed (eating, walking, grooming) at which point the mouse was returned to IVC cage housing.

The mice underwent µCT scanning on the day of surgery and at 2, 4, 6, and 8 weeks post-surgery. Scanning at the 8-week time point was performed post-mortem and included *ex vivo* hindlimbs (**Supplementary Table 2**). μCT reconstructions were obtained using MILabs software (MILabs-Recon v. 11.00) at 40 µM and 20 µM resolution for *in vivo* and *ex vivo* images respectively. Formation of skeletal tissue was assessed using Imalytics Preclinical software v3.0 (Gremse-IT GmbH) and a gauss filter of 1.0 was applied in each method of analysis. A bone density phantom was scanned at each time point using identical parameters as a reference for bone density quantification to enable a minimum threshold to be set for bone volume analysis.

For analysis, segmentation of the region of interest was performed at all timepoints. Initially analysis of a segment of bone from proximal to distal femur was quantified for an overall view of bone formation in each group (**Supplementary Figure 10)**. Further focussed analysis was performed by selecting a 4 mm diameter 6 mm length cylindrical volume over the scaffold (scaffold size 3 mm diameter × 5 mm length), therefore including a 0.5 mm periphery around scaffold. The centre of the scaffold was measured to be 2.5 mm from the femur ends and 1.5 mm from the perceived edge of the scaffold. This was most accurate in *ex vivo* images due to enhanced detail of the radiolucent scaffold against the surrounding soft tissues (**Supplementary Figure 11**). The cylindrical volume was applied to the scaffold (**Supplementary Figure 12 A).** For collagen sponge, the 4 mm diameter × 6 mm length cylinder was used with 0.5 mm of both sides of the femur included in analysis as this was equivalent to the limbs with the scaffolds to avoid underestimation of bone volume. Nevertheless, the bone volume would be difficult to equivocate to the PCL-TMA900 scaffolds as the diameter of the collagen sponge swelled to greater than 4 mm, so the outer bone was disregarded from quantification (**Supplementary Figure 12 B**).

To analyse the volume of bone formed on the scaffold a 3 mm diameter × 5 mm length cylinder was selected, and bone volume quantified in the *ex vivo* images at week 8, permitting enhanced visualisation of the shape of bone formed within the scaffold (**Supplementary Figure 13 A)**. In the case of the collagen sponge, the 3 mm × 5 mm cylindrical volume was selected, and the femur ends removed from the bone quantification as the sponge was compressed to bring the femur ends within the 3 mm × 5mm volume, which would lead to overestimation of bone volume. Again, the outer bone which formed wider than 3 mm diameter was disregarded from analysis (**Supplementary Figure 13 B**).

The medial and lateral sides of the most efficacious scaffolds; ELP/PEA/FN/BMP-2, PEA/FN/BMP-2 and Laponite/BMP-2 coated scaffolds were quantified by creating a cylinder (1.5 mm × 5 mm) medial and lateral within the 3 mm × 5mm scaffold cylinder volume (**Supplementary Figure 14 A-F)**. The cylinder was positioned over the medial and lateral scaffold respectively by lining up the image in all 2D planes. The medial and lateral volumes were segmented out and quantified individually (**Supplementary Figure 14 G and H)**.

### Histological processing, embedding, and sectioning of mouse limbs

The polyimide IM pin was not removed from each limb as the IM pin was amenable to processing, embedding and sectioning (**Supplementary Figure 15**). Samples were fixed in 4% PFA for 3-4 days rinsed in PBS and decalcified for 4 weeks with 5% ethylenediaminetetraacetic acid (EDTA) in 0.1 M Tris in dH_2_O (pH 7.3 with NaOH) at 4°C on the rotator. The decalcification solution was changed weekly, and µCT/x-ray confirmed complete decalcification prior to rinsing in PBS and dehydration. After dehydration through ethanol solutions (70%, 90%, 100% twice) and chloroform:100% ethanol mix in a 1:1 ratio, followed by 100% chloroform twice, for 1 hour in each solution, the samples were processed in the Heraeus Vacutherm VT6025 Vacuum Oven in molten paraffin wax (Histowax, Leica) at 65°C for 1 hour. Fresh wax was added and a further 2 hours in the vacuum oven commenced. The samples were embedded in fresh molten wax and cooled at 0-1°C for 2 hours before storage at 4°C. The blocks were sectioned at 10 µM on a Microm 330 microtome (Optec, UK). The sections were transferred via a water bath to pre-heated glass slides for 2 hours until dry and placed in the slide oven at 37°C for 4-6 hours prior to storage at 4°C.

### Histological staining of tissues

#### Alcian blue/Sirius red

The method was extrapolated from Marshall *et al*. [9]. The tissue sections on slides were rehydrated through Histoclear (2 × 7 minutes), ethanol solutions of 100% (twice) to 90% to 50% (2 minutes each), followed by immersion in water. Weigert’s Haematoxylin was applied for 10 minutes, removed by immersion 3 times in acid/alcohol (5% HCl/70% ethanol) followed by washing in the water bath for 5 minutes. Slides were immersed in 0.5% Alcian blue 8GX in 1% acetic acid for 10 minutes, 1% molybdophosphoric acid for 10 minutes, followed by rinsing with water prior to staining with 1% Picrosirius Red (Sirius Red) for 1 hour. Excess stain was rinsed off in water and the slides were dehydrated in increasing concentrations of ethanol of 50%, 90%, 100% twice and Histoclear twice for 30 seconds in each, prior to mounting with dibutyl phthalate xylene (DPX) and a glass coverslip and air-dried.

#### Goldner’s Trichrome

The method was extrapolated from Marshall *et al.* [9]. The method followed is identical to Alcian blue/Sirius Red staining to the stage of acid/alcohol immersion and washing in water for 5 minutes. Ponceau Acid Fuchsin/Azophloxin (Sigma) was applied for 5 minutes followed by a 15 second wash in 1% acetic acid. Phosphomolybdic acid/Orange G was applied for 20 minutes followed by another 15 second wash in 1% acetic acid. Light Green was applied for 5 minutes followed by the third 15 second wash in 1% acetic acid. The sections on the slides were dehydrated in ethanol 90% and 100% twice and Histoclear twice for 30 seconds each prior to mounting with DPX and a glass coverslip and air-dried.

#### Tartrate-resistant acid phosphatase (TRAP) staining

For the visualisation of osteoclast activity, TRAP Basic Incubation Medium (9.2 g sodium acetate anhydrous, 11.4 g L-tartaric Acid, 950 mL dH_2_O, 2.8 mL glacial acetic acid, pH adjusted with 5 M sodium hydroxide to pH 4.7-5.0 and made up to 1 L with dH_2_O and Naphthol AS-MX Phosphate substrate (2 mg Naphthol AS-MX Phosphate to 0.1 mL of ethylene glycol monoethyl ether) were prepared. The final staining solution was prepared from 20 mL of TRAP basic incubation medium, 0.1 mL of Naphthol AS-MX Phosphate substrate and 12 mg of Fast Red Violet LB Salt, prewarmed to 37°C in a drying oven. The slides were dewaxed and rehydrated prior to staining with TRAP staining solution applied to each slide and incubated at 37°C for 30 minutes, checking red positive stain development on positive control slides. The slides were rinsed in dH_2_O and counterstained with 0.02% Fast Green stain for 30 seconds, rinsed in dH_2_O, dehydrated through 50%, 90%, 100% (twice) and Histoclear (twice) for 30 seconds in each solution. Slides were mounted with DPX and a glass cover slip applied and allowed to dry.

#### Image acquisition of histology slides

Histological wax samples were imaged using the Zeiss Axiovert 200 microscope using bright field microscopy and images captured with the Axiovision 4.2 imaging software (Zeiss). Resin embedded samples were imaged using a Leica TCS-SP8 microscope with dual excitation at 405 nm and 458 nm and primary detection limits of 480-560 nm and red filter was set at 600-690 nm. The Alizarin red stained resin embedded tissues were imaged using an Epson Perfection V700 Photo Dual Lens System scanner.

### Statistical analysis

Data from replicates were analysed using on GraphPad Prism 9, version 9.2.0. P values <0.05 were considered significant. Graphical representation of significance as follows: *p<0.05, **p<0.01, ***p<0.001, ****p<0.0001. All data presented as mean and standard deviation (S.D.).

## Results

### Gross dissection

There were no signs of inflammation or infection around any of the constructs examined during the study. In mice with the PCL-TMA900 scaffolds implanted, limb movement was restricted in abduction and full retraction, however, the mice would not usually perform such movements when conscious, therefore this did not hinder normal activity or appear to cause discomfort. The scaffold was firmly held abutting the proximal femur; therefore, the distal edge of the scaffold was palpable and the distal femur adducted inwards, due to the pull of the quadriceps and contraction of the medial soft tissues. This caused an almost 90° bend in the plastic pin at the proximal end of the distal femur fragment (**Supplementary Figure 16**).

### µCT analysis of bone formation within the femur segment and scaffold region

Initially on the day of surgery, the transected limbs were not aligned in all planes, as the pin was not tight within the marrow cavity allowing some pin movement. Due to a conservative approach adopted in the ostectomy surgery, the operative limb was longer than the contralateral limb initially as the bone fragment removed was smaller (approximately 3 mm in length) than the 5 mm long scaffold (**Supplementary Figure 17**). The distal limbs displayed a clear angulation at week 2 seen on µCT due to bending of the plastic pin at the proximal extent of the distal femur (**Supplementary Figure 18**). The limbs were stable as the soft tissues contracted medially and thus the defect was not able to be manipulated or destabilised from week 2 onwards as observed while mice were anaesthetised for µCT image acquisition. The mice with ELP/PEA/FN/BMP-2 coated scaffolds displayed more aligned limbs, although the method of surgery was the same. Cranial angulation of the limb was apparent in the ELP/PEA/FN/BMP-2 coated scaffold group, rather than medial bending (**Supplementary Figure 19**). The collagen sponge and Laponite/BMP-2 coated scaffolds displayed a marked bone formation response from week 2 onwards with bone formation bridging the defect over the 8-week study (**Supplementary Figure 18 and 19**). The bone formed within a segment from the neck of femur to distal femur was quantified however, the medial angulation led to variation in bone formation (**Supplementary Figure 20).** The average bone volume of each group was comparable at the start of the study. At week 2, a marked increase in the bone volume of the collagen/BMP-2 group was observed. There was a subsequent reduction in volume of the collagen/BMP-2 group over the following 6 weeks. The remaining groups displayed a temporal increase in bone volume to week 4 before a plateau was observed until week 8, with limited progression in bone formation (**Supplementary Figure 20**). On further analysis, the average bone volume for each group displayed a similar trend of bone formation to week 4 and no further obvious progression. The collagen/BMP-2 bone volume was observed to increase to week 2 and to subsequently decrease in bone volume, likely due to bone remodelling (**Supplementary Figure 21**). When analysed as a percentage increase in bone volume, there was no significant difference between all the groups examined (**Supplementary Figure 22**).

At week 8, the µCT scans displayed union of the femur ends via the PCL-TMA900 scaffold in the Laponite/BMP-2 coated PCL-TMA900 scaffold group, and via a bulbous callous in the collagen sponge/BMP-2 group. There was no union of the fracture in the ELP, Laponite, PEA/FN/BMP-2, ELP/PEA/FN/BMP-2 coated PCL-TMA900 scaffold groups and the uncoated scaffold served as the comparator with no obvious bone formation over or within the uncoated PCL-TMA900 scaffold (**Figure 1**).

**Figure 1:**
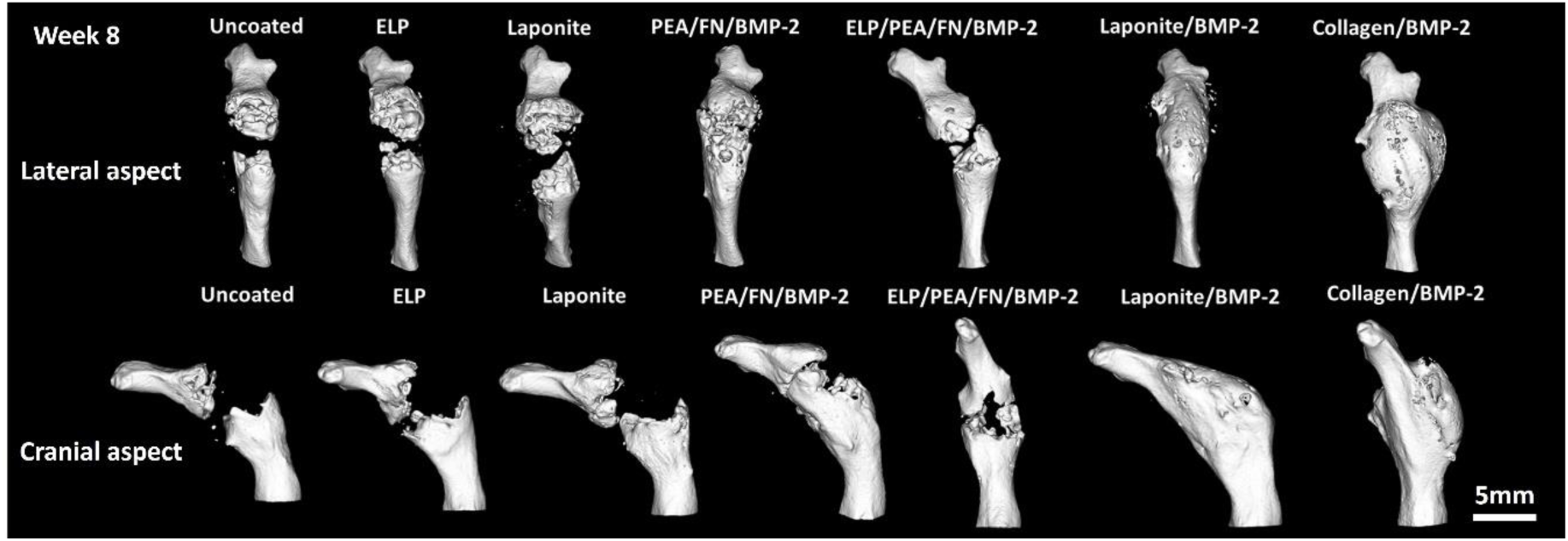
Representative µCT images from the lateral and cranial aspects of the limb with maximal bone formed in each group. Due to the pin bending, an inward, medial bend to the limb resulted in all mice except the ELP/PEA/FN/BMP-2 group. Over time the Laponite/BMP-2 coating displayed the largest bone healing compared to the other coating groups. The collagen/BMP-2 group displayed complete union of the fracture site with smooth remodelling of the cortex craniomedially and a spherical shell of bone laterally and caudally. Representative images shown from each group (n=6, n=5 for ELP/PEA/FN/BMP-2), scale bar 5 mm.

The *ex vivo* µCT scans were comparable to the *in vivo* scans with the union of the fracture in the Laponite/BMP-2 and Collagen sponge/BMP-2 groups only (**Supplementary Figure 23**).

Bone volume was analysed using a 6 mm × 4 mm cylindrical volume around the defect site. A significant difference was found when comparing the uncoated PCL-TMA900 to the Laponite/BMP-2 and collagen/BMP-2 constructs (**Figure 2 A).** Additionally, when analysis focussed on the 3 mm × 5 mm cylindrical volume of the scaffold itself or collagen sponge constructs to determine the bone formed within the scaffold volume, there remained a significant difference between the uncoated PCL-TMA900 scaffold and the Laponite/BMP-2 and collagen/BMP-2 groups (**Figure 2 B**). Further, the medial and lateral aspects of the 3 mm × 5 mm cylindrical volume were analysed demonstrating the Laponite/BMP-2 coated PCL-TMA900 scaffold contained significantly greater bone formation laterally than the PEA/FN/BMP-2 coated PCL-TMA900 scaffold. There was no significant difference on the medial aspect between the three scaffold coating groups (**Figure 2 C**).

**Figure 2:**
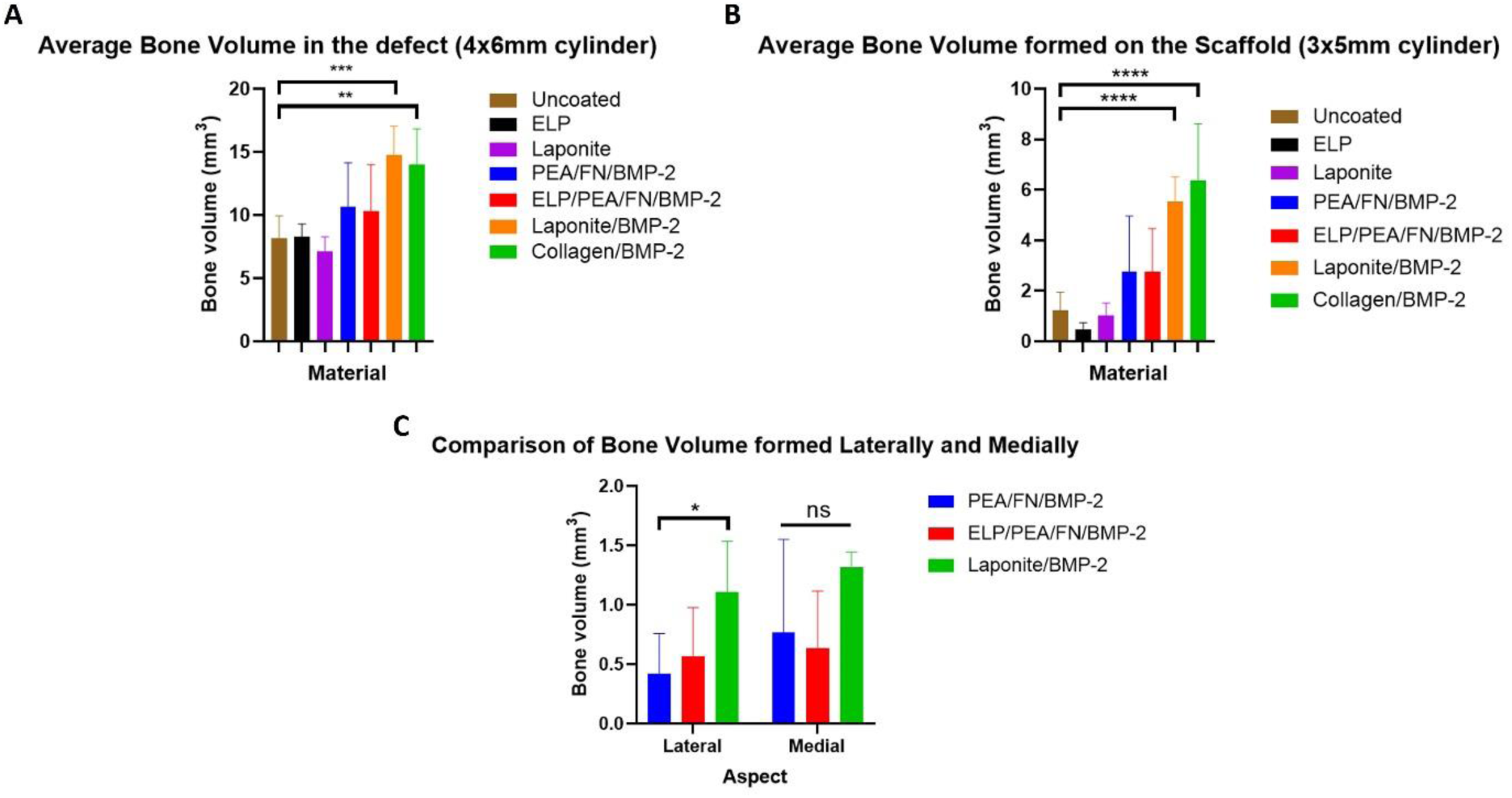
Quantification of bone volume using *ex vivo* images at week 8. (A) Quantification of a 4 mm × 6 mm cylinder revealed the Laponite/BMP-2 and collagen sponge/BMP-2 constructs had significantly greater bone formation than the uncoated PCL-TMA900 scaffolds, while all other groups did not display a significant difference to the uncoated PCL-TMA900 scaffolds. N=6 each group, except n=5 for ELP/PEA/FN/BMP-2 group, one-way ANOVA with Dunnett’s multiple comparisons test, **p<0.01, ***p<0.001, mean and S.D. shown. (B) Quantification of bone volume in a 3 mm × 5 mm cylinder confirmed that only the Laponite/BMP-2 and collagen sponge/BMP-2 constructs displayed significantly greater bone formation than the uncoated PCL-TMA900 scaffolds. N=6 each group, except n=5 for ELP/PEA/FN/BMP-2 group, one-way ANOVA with Dunnett’s multiple comparisons test, ****p<0.0001, mean and S.D. shown. (C) Medial versus lateral bone formation in a 3 mm × 5 mm cylinder volume of the scaffold. A significant difference was found between the lateral aspect of the scaffold in Laponite/BMP-2 coated and PEA/FN/BMP-2 coated groups. There was no significant difference on the medial aspect of the scaffold between any of the 3 groups. N=6 each group, except n=5 for ELP/PEA/FN/BMP-2 group, 2-way ANOVA with Tukey’s multiple comparisons test performed, ns, not significant, *p<0.05, mean and S.D. shown.

### Histology demonstrated intramembranous and endochondral ossification during bone formation

Histology of the limbs indicated distinct populations of cells within the tissues as the bone healing process progressed. The limbs selected for histology images were those with the greatest degree of bone formation on µCT results to enable bone formation analysis if present upon sectioning. The angulation of the limbs and scaffold within each defect can be seen (**Supplementary Figure 24**), with bone formation as per the µCT images. The bone shelf formed medially by intramembranous ossification from the periosteum which abutted the scaffold. Chondrocytes could be observed, illustrating endochondral ossification in action on the medial aspect of the limb. The lateral aspect of the uncoated scaffold was covered in fibrous tissue (**Figure 3 A**). The ELP coated PCL-TMA900 scaffold displayed similar findings with new bone formation mostly medially and chondrocytes extending from this bone proximally along the medial aspect of the scaffold, extending towards the proximal extent of the femur (**Figure 3 B**). This was likely due to the exaggerated angulation of the limb and biomechanics encouraging the medial bone to form more than on the lateral aspect of the limb. The Laponite only coated PCL-TMA900 scaffolds encouraged no bone to form along or within the scaffold, with only the bone tissue forming on the medial aspect of the limb, again likely due to the angulation and forces on the limb, similar to the uncoated and ELP only coated scaffolds, which also did not induce endochondral ossification or bone formation around or within the scaffold (**Figure 3 C**). In the PEA/FN/BMP-2 coated PCL-TMA900 scaffold, the tissues had infiltrated into the pores of the scaffold and ossified, producing cortical type bone with regularly spaced osteocytes surrounding marrow cavities of cancellous type bone, which was vascularised, as evidenced by blood cells seen in the Goldner’s trichrome stained sections (**Figure 3 D**). Bone was observed to be formed predominantly medially and dorsally in this example mouse. The ELP/PEA/FN/BMP-2 coated scaffold had minimal bone formed in the centre of the scaffold on µCT analysis although, bone tissue extending from proximally and distally around the periphery of the scaffold could be seen. The bone was mature cortical bone abutting fibrous tissue which extended into the pores of the scaffold, with chondrocytes gathered here as endochondral ossification was progressing (**Figure 3 E**). The Laponite/BMP-2 coated PCL-TMA900 scaffolds showed the most pronounced and uniform new bone formation on µCT and this was confirmed upon histological staining. The periphery of the scaffold was encased in cortical bone with osteocytes within the matrix and areas of osteoid as the bone matured, while the pores of the scaffold were lined with cortical bone surrounding bone marrow, with blood cells seen within the marrow cavities (**Figure 3 F**). The collagen sponge with BMP-2 formed smooth cortical bone craniomedially and a thin spherical shell caudolaterally which was seen by µCT and grossly upon dissection. The histology showed thicker cortical bone bridging the distal femur to proximal femur medially, with more fragile, less directionally orientated bone laterally, as noted on the µCT images (**Figure 3 G**).

**Figure 3:**
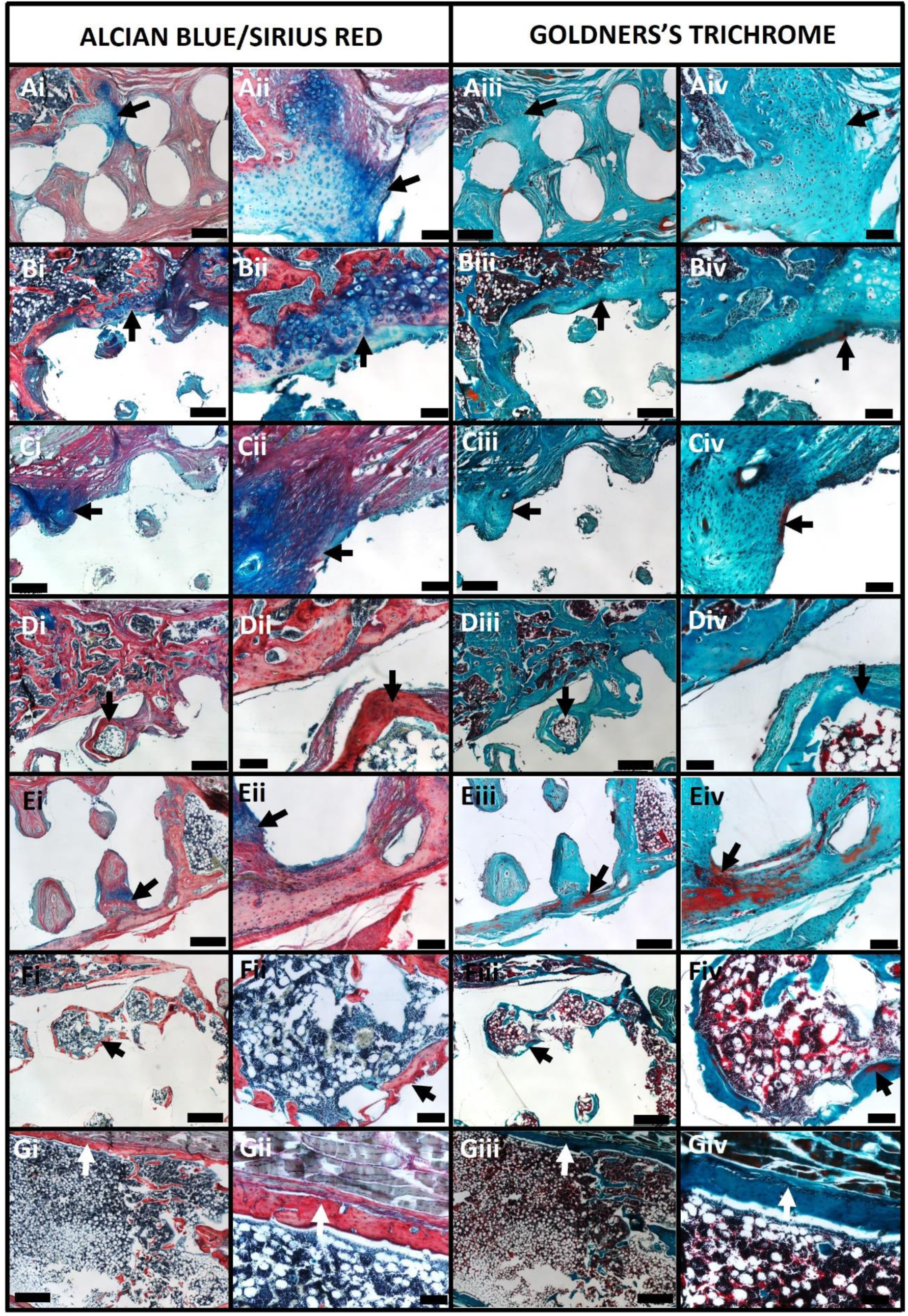
Histological analysis of bone formation surrounding the PCL-TMA900 scaffolds and collagen sponge/BMP-2 construct using Alcian Blue/Sirius Red (AB/SR) and Goldner’s Trichrome (GT) staining. The medial aspect of the limb is to the top and lateral aspect to the bottom, proximal femur to the right and distal femur to the left of each image. (A) Uncoated PCL-TMA900 scaffold: (Ai, Aii) A population of chondrocytes in blue-staining proteoglycan matrix around the structure of the scaffold medially (black arrow). (Aiii, Aiv) The medial new bone and a population of chondrocytes in within tissue surrounding the structure of the scaffold medially (black arrow). (B) ELP coated PCL-TMA900 scaffold: (Bi, Bii) The new bone medially and chondrocytes clusters (black arrow) at the junction between the new bone and the scaffold itself with fibrous tissue within the cylindrical pores of the scaffold. (Biii, Biv) Osteoid tissue (black arrow) forming along the scaffold medially at the edge of the chondrocytes. (C) Laponite coated PCL-TMA900 scaffold: (Ci, Cii) Red fibrous collagenous tissue with a large area of blue proteoglycan matrix and chondrocytes on the medial aspect of the limb forming a thin area of bone matrix (black arrow). (Ciii, Civ) Osteoid tissue (black arrow) surrounded by proteoglycan-rich matrix and cellular environment. (D) PEA/FN/BMP-2 coated PCL-TMA900 scaffold: (Di, Dii) The bone medially with islands of marrow cavity within a layer of cortical bone formed around the coated scaffold material. Bone tissue formed within the pores of the scaffold as cylindrical areas of cortical bone enclosing a central marrow cavity (black arrow). (Diii, Div) The new mature cortical bone along the edge of a pore within the scaffold (black arrow). (E) ELP/PEA/FN/BMP-2 coated PCL-TMA900 scaffold: (Ei, Eii) Mature cortical bone with osteocytes embedded in the calcified matrix while fibrous tissue surrounded the scaffold and within the pores with differentiation of chondrocytes potentially leading to further bone tissue formation (black arrow). (Eiii, Eiv) The osteoid within the cortical bone (red arrow). (F) Laponite/BMP-2 coated PCL-TMA900 scaffold: (Fi, Fii) Cortical bone within the pores of the scaffold (black arrow) and marrow cavities within the pore spaces. (Fiii, Fiv) The osteoid tissue (black arrow) formed, with extensive red blood cells seen within the marrow cavity enclosed by the new bone. (G) Collagen/BMP-2 construct: (Gi, Gii) The cranial aspect of the collagen sponge/BMP-2 construct showed the medial bone was uniform and over 100 µm thick in regions (white arrow) with a large marrow cavity formed within the bone shell. (Giii, Giv) GT staining, with the cortical bone (white arrow) and vascular marrow cavity seen. (i and iii columns scale bar 500 µm, ii and iv columns scale bar 100 µm).

TRAP staining was used to visualise osteoclast location to assess regions of remodelling. The histology showed osteoclasts were active around the scaffold and at the proximal and distal bone ends. Osteoclasts were observed within the pores of the scaffolds and uniformly along the edge of the bone formed in the collagen sponge/BMP-2 group (**Figure 4**).

**Figure 4:**
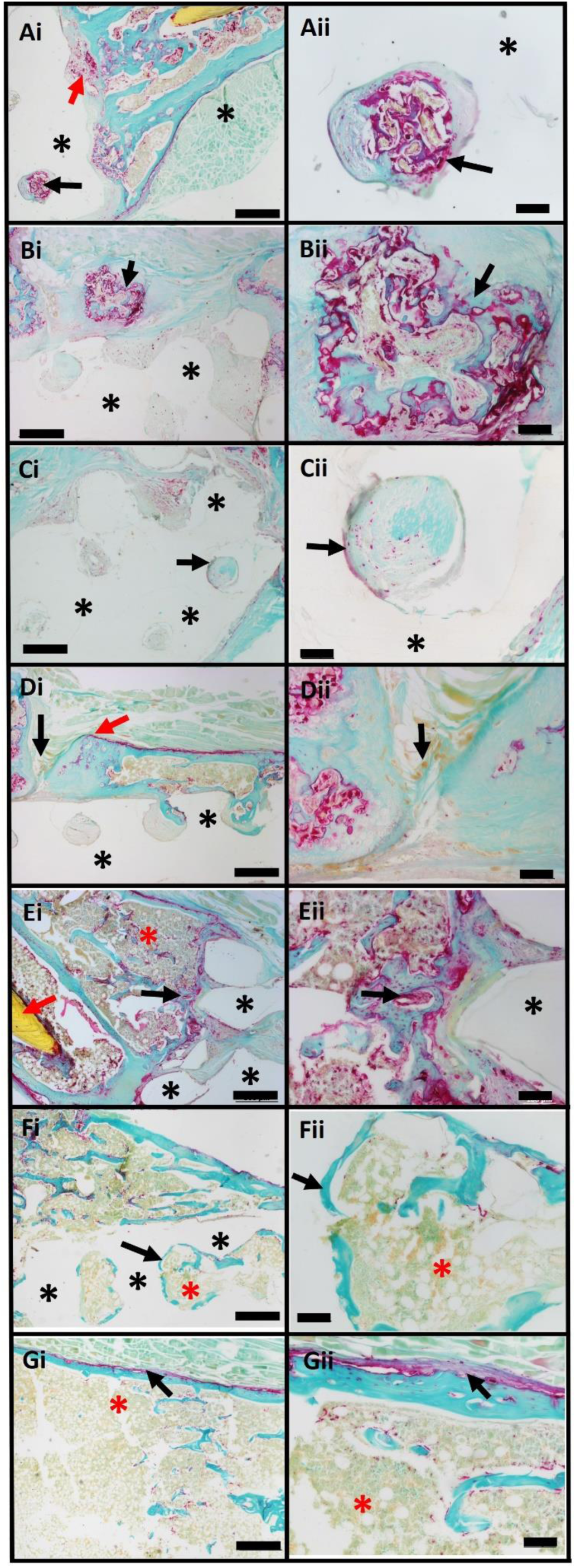
TRAP staining of histological sections to visualise regions of osteoclast activity. (Ai, Aii) Uncoated scaffold; osteoclasts at the junction between the scaffold (*) and proximal end of the femur (red arrow) and within a pore of the scaffold (black arrow). (Bi, Bii) ELP coated scaffold; osteoclasts (black arrow) around a mass of bone on the medial aspect of the limb adjacent to the scaffold (*). (Ci, Cii) Laponite coated scaffold; relatively fewer osteoclasts were seen around the scaffold (*), with some within the pores of the scaffold. (Di, Dii) PEA/FN/BMP-2 coated scaffold; osteoclasts (red arrow) lining the medial bone surface over the scaffold (*), and an area where bone from the proximal and distal femur has almost united (black arrow). (Ei, Eii) ELP/PEA/FN/BMP-2 coated scaffold; osteoclasts at the junction between the new bone and the scaffold (*) (black arrow), with new bone formed by intramembranous ossification from the periosteum seen (red asterix) and the plastic pin (red arrow) seen within the section. (Fi, Fii) Laponite/BMP-2 coated scaffold; quiescent cortical bone (black arrow) and mature marrow cavities (red asterix) formed within the pores of the scaffold material (*), with less osteoclast staining seen. (Gi, Gii) Collagen/BMP-2; osteoclasts (black arrow) lining the new cortical bone around the marrow cavity (red asterix) formed by the collagen sponge/BMP-2 construct. (Left column scale bar 500 µm and right column scale bar 100 µm).

## Discussion

This study set out to apply 3D printed PCL-TMA900 scaffolds with various bioactive coatings in a critical-sized femur defect model to assess bone formation compared to uncoated PCL-TMA900 scaffolds in an orthotopic site. Laponite with BMP-2 as a coating on the PCL-TMA900 scaffold material, was successful in uniting the fracture with significant bone formation around and within the porous scaffold. Non-union of the bone ends in mice with the uncoated PCL-TMA900 scaffold and bridging bone formation using the collagen sponge with BMP-2 (5 µg) was observed, confirming suitable scaffold length, scaffold material and mass of BMP-2 applied to the positive control.

The current study set out to examine the different coating options (ELP, PEA/FN/BMP-2, ELP/PEA/FN/BMP-2 or Laponite/BMP-2) on the PCL-TMA900 scaffolds to determine which was most efficacious to go forward to the large animal bone defect model. In the study, the Laponite/BMP-2 coated scaffolds were the only coated scaffold group to show a markedly significant difference from the uncoated PCL-TMA900 in bone volume formed over the 8-week study. Therefore, the hypothesis that coating a scaffold with an osteoinductive coating would significantly improve bone formation was confirmed. The Laponite/BMP-2 coating on PCL-TMA900 united the fractured bone ends and contained new bone at the periphery of the scaffold and within the scaffold pores, indicating adequate scaffold pore size with no surplus, ectopic bone formation in the surrounding soft tissues. The Laponite/BMP-2 coating was used immediately once dry as previous work has shown that a delay in use may affect efficacy [8].

ELP coating on a solid scaffold material has been shown to enhance integration between the scaffold and bone but had not been found to significantly contribute to bone formation when compared to uncoated controls [12]. The PEA/FN/BMP-2 coating has previously been applied with successful bone formation in murine radial defects and veterinary clinical cases, when combined with autograft [13, 34]. However, it was not found to produce reliable, consistent bone volumes in this study, potentially as a consequence of the PEA/FN/BMP-2 coating relying on a greater surface area such as on porous granular materials to deliver a critical mass of BMP-2 to take advantage of the PEA/FN/BMP-2 reciprocal binding. The ‘critical mass’ of BMP-2 to induce bone formation in rodents has been deemed as ∼1 µg of BMP-2, but a lower mass may be efficacious if sequestered in the bone fracture site to provide a consistent stimulus for osteogenic differentiation of skeletal cell populations [16]. Laponite’s mechanism of action also relies on this principle of sequestering BMP-2 to provide bone formation using lower masses of BMP-2, reducing cost and potential adverse effects clinically.

Consistently, the lack of mineralisation around the uncoated PCL in the first pilot study or PCL-TMA900 in the second pilot study and current study was unsurprising, given the absence of an osteogenic component on the scaffold surface to induce bone formation. This critically, highlights the requirement for an *in vivo* validated osteogenic component on 3D printed scaffolds to enhance their potential as biodegradable scaffolds for bone augmentation. The collagen sponge delivering 5 µg of BMP-2 was sufficient as a positive control. The bone bridging induced from the collagen sponge began some distance from the transected bone ends as the transected bone ends did not ‘regrow’ themselves. BMP-2 can affect the periosteum and endosteum to recruit progenitor cells to differentiate towards chondrogenic or osteogenic lineages respectively [35]. Therefore, bone forms as an outer shell from the periphery of the sponge with the periosteum appearing to play a significant role in forming a bone shell by intramembranous ossification.

One mouse in the ELP/PEA/FN/BMP-2 coated group died on day 19 post-operatively due to a systemic, presumably anaerobic, bacterial infection. There were no signs of lameness, inflammation, or bone lysis present at the week 2 µCT analysis, therefore the infection was possibly unrelated to the study. At week 8, mouse 4 in the uncoated PCL-TMA900 group was noted to have 2 cavities within the proximal femur, likely due to pressure resorption due to the plastic pin against the bone, however, the final bone volume of the segment was comparable to the other 5 mice in this group and thus was included in the analysis of the femur segment volumes but regardless, the quantification of bone volume around the scaffold would not have been affected by this finding.

Previous reports describe stabilising the fractured femur with an intramedullary (IM) needle from the greater trochanter through the distal femur and stifle joint [27, 36, 37]. This published method ensured adequate stability of the fracture site and material held within the bone defect, however it was not conducive to sequential, quantitative radiographic imaging. Clough and colleagues inserted a metal pin via the transected bone ends with a metal spacer to maintain the gap in the bone, which was radiodense [33]. Removal of the metal spacer implant was described by sharp dissection prior to histology which would potentially damage the delicate forming tissue, while the use of a polyimide plastic pin negates this issue [33]. The IM pin insertion method chosen was to limit surgical trauma to the femur and prevent decimation of the entire bone marrow cavity. Consideration was given for thermal necrosis and osteoclastic activity at the bone ends due to the bone saw, however, this bone lysis was not observed to the same degree when the plastic pin was used compared to the metal pin, perhaps due to less stress shielding of the surrounding cortical bone. In the first pilot study using metal IM pinning, one mouse suffered a complication of displacement of the distal femur segment at week 8, likely due to the density of the pin protecting the scaffold and surrounding bone from stimulatory forces, leading to resorption of bone in the distal femur segment over the preceding weeks.

An attempt to ameliorate the scatter and beam hardening artefacts using a copper filter during µCT scanning was made, but there remained artefacts from the metal. The quantification of bone volume when using the plastic pin was thought to be markedly more accurate, with reduction in human error compared to the necessity of manual selection of areas to disregard due to scatter from the metal pin, when using the analysis software [33]. The mice were scanned every 2 weeks to monitor bone formation over time. No detrimental effect on healing of the fracture callus has been found with sequential weekly µCT scans, and the inclusion of control mice allowed comparison of data generated [38].

The plastic pin could be used to stabilise the bone ends and alleviate difficulties in µCT imaging and histology associated with use of metal stabilisation. Results showed that the plastic pin was successful in holding the scaffold in position between the bone ends, permitted normal locomotion despite bending, could be imaged by µCT, and sectioned for histological analysis. This pilot study also crucially confirmed the biocompatibility of the PCL-TMA900 material scaffolds, with no adverse effects noted on gross or histological examination, similar to a previous subcutaneous implantation study [8]. Greater bone formation in the plastic IM pin group compared to the metal IM pin was found, however, this result was not statistically significant.

The PCL-TMA900 scaffolds and plastic pin were amenable to processing and sectioning during histological evaluation of bone formation. The histology by Kaur *et al.* mirrors the findings of the current studies with intramembranous bone formation between the periosteum and cortex of the transected bone ends [39]. It was found that the compressive strain medially promoted the formation of cartilaginous tissue, promoted by the cells at the edge of the transected end of the femur. The tensile strains laterally induced the formation of fibrous tissue across the scaffolds, however, the Laponite/BMP-2 and collagen/BMP-2 constructs had mature mineralised bone both medially and laterally surrounding each construct, as the scaffold delivery of BMP-2 induced bone formation equally around the scaffold while this did not occur in the other scaffold coating groups [40]. The TRAP staining indicated osteoclast activity around the scaffolds and within the pores of the scaffolds in each group, however, the Laponite and Laponite/BMP-2 groups appeared to have less osteoclast activity within the pores of the scaffold, due to predominantly collagenous tissue in the Laponite only group and the maturity of the bone formed in the Laponite/BMP-2 group with organised bone surrounding marrow cavities within the pores of the scaffold.

The mice had no rigid fixation of the scaffold within the fracture defect and could weight bear immediately postoperatively. The plastic pin fixation method was unstable initially with potential movement at the scaffold/bone end interface, but the soft tissues held the scaffold firmly between the bone ends by week 2. The micromotion induced healing by endochondral ossification, with significant numbers of chondrocytes seen in histological sections around the distal bone end medially and on the periphery of the scaffold. The strain upon the fracture fragments, or in this case the scaffold and transected femur, were found to have an optimal level for healing. Positive correlations have been found between fracture activity in the early post operative 1-2 week period with low strain leading to direct primary healing, moderate strain leading to secondary healing via callus formation, with the motion reducing as healing progresses [41]. However, too large a strain leads to cessation of healing and non-union of the fragments/construct and implant failure [41, 42]. Conversely, if the fixation method is too rigid, healing will be abated leading to the indication that there is an optimal strain in a fracture site to initiate and propagate healing [40]. Cyclic compression has been found to lead to RUNX2 and BMP-2 production or increased sensitivity to BMP-2 in cells, leading to osteogenic differentiation [43]. The current model shows inverse dynamization, that is the flexible fixation becomes more rigid in the first 2 weeks in that as the limbs curved, the scaffolds prevented movement of the distal femur, leading to a more stable environment for further healing [44]. The stress or load would have been applied in the mice by weight bearing, especially when standing on their hindlimbs or through muscle contraction [40].

Poly(caprolactone) trimethacrylate (PCL-TMA900) was chosen as a material which would biodegrade due to the presence of ester bonds and because of the material properties suitable for SLA 3D printing (i.e., lower glass transition temperature). The molecular weight was chosen as a compromise between suitable material properties (i.e., toughness and stiffness), whilst having a viscosity suitable for 3D printing. In an ideal situation, the 3D printed scaffold material would degrade on a timescale shortly after the formation of new bone in and around the scaffold, to allow for complete healing of the defect. While this is not the case within the present study as the scaffolds are still present after 8 weeks, due to their chemical composition, it would be expected that the scaffolds would degrade on a larger time scale (i.e., months to years). The complex environment into which the scaffolds are implanted makes correlation of *in vitro* degradation or accelerated degradation conditions to *in vivo* degradation challenging. Ultimately, longer follow up studies would be required to verify and tailor the degradation times. Crucially, we have demonstrated that the 3D printed PCL-TMA900 scaffolds show excellent biocompatibility and can support the formation of new bone with a suitable osteogenic coating.

Due to the significant bone formation surrounding the Laponite/BMP-2 coated PCL-TMA900 scaffolds uniting the bone ends in this study, further studies with Laponite/BMP-2 coating in the ovine femoral condyle defect model is warranted to scale up the Laponite/BMP-2 coated construct to assess repeated efficacy of the Laponite/BMP-2 coating on PCL-TMA900 scaffolds prior to clinical translation.

## Conclusion

This study used a novel murine femur bone defect method to confirm the biocompatibility and efficacy of Laponite/BMP-2 as a bioactive coating on rigid PCL-TMA900 polymer scaffolds for bone repair. The Laponite/BMP-2 coating induced solid union of the fracture due to significant mature bone formation on and within the PCL-TMA900 scaffold as seen by sequential µCT imaging and histology. This Laponite/BMP-2 coating on PCL-TMA900 scaffolds will be taken forward in an ovine femoral condyle defect model, to investigate the ability of the BMP-2 to induce bone formation when the scaffold shape is developed with an increase in magnitude of the bone defect in a critical-sized ovine metaphyseal bone defect. The ovine model will provide a necessary test of osteogenic efficacy and scaffold material stability on the path towards clinical application of the Laponite/BMP-2 coated PCL-TMA900 material for bone repair.

## Credit author statement

Callens, S. J. P. designed and modelled the scaffolds. Echalier, C. and Wojciechowski, J. P. produced the PCL-TMA900 resin. Wojciechowski, J. P. and Yang, T. prepared and printed the PCL-TMA900 scaffolds. Jayawarna, V. extrusion printed the PCL scaffolds for the first pilot study, performed PEA coating of scaffolds and organised the EO sterilisation of materials. Hasan, A. ELP coated the scaffolds. Dawson J. I supplied the Laponite and edited the paper. Marshall K. M. conceptualisation of the study methods and conducted the *in vivo* experiments, data collection and analysis, histology, and wrote the paper. Kanczler J. M., Mata, A., Salmeron-Sanchez, M., Stevens, M. M., and Oreffo R. O. C. were responsible for conceptualisation, funding acquisition, study supervision and editing of the manuscript.

## Data and materials availability

All data associated with this study are presented in the paper or the Supplementary Materials. All raw data is available upon request.

## Acknowledgements

Research support for this study from the Biotechnology and Biological Sciences Research Council (BBSRC BB/P017711/1), the UK Regenerative Medicine Platform Acellular / Smart Materials – 3D Architecture (MR/R015651/1), ERC Proof-of-concept grant MINGRAFT, and University of Southampton is gratefully acknowledged as well as many useful discussions with current members of the Bone and Joint Research Group in Southampton, UK. We thank Professor Nicholas Evans, University of Southampton, for helpful discussions on the programme of work. Dr Katie Dexter, University of Southampton Biomedical Imaging Unit, is acknowledged for advice in µCT scanning of mice and thanks to Professor Nicholas Evans, Dr Roxanna Ramnarine Sanchez and Mr Andrew Rawlings, at the University of Southampton, for their assistance during the final *in vivo* study.

## Appendix A. Supplementary data

Supplementary data to this article can be found online.

## References

1. Fernandez de Grado, G., et al., Bone substitutes: a review of their characteristics, clinical use, and perspectives for large bone defects management. J Tissue Eng, 2018. 9: p. 2041731418776819.

2. Keating, J.F., A.H.R.W. Simpson, and C.M. Robinson, The management of fractures with bone loss. The Journal of Bone and Joint Surgery, 2005. 87(2): p. 142–150.

3. Shegarfi, H. and O. Reikeras, Review article: Bone transplantation and immune response. Journal of Orthopaedic Surgery, 2009. 17(**2**): p. 206–211.

4. Haugen, H.J., et al., Bone grafts: which is the ideal biomaterial? J Clin Periodontol, 2019. 46 **Suppl 21**: p. 92–102.

5. Amini, A.R., C.T. Laurencin, and S.P. Nukavarapu, Bone tissue engineering: recent advances and challenges. Crit Rev Biomed Eng, 2012. 40(5): p. 363–408.

6. Bahney, C.S., et al., The multifaceted role of the vasculature in endochondral fracture repair. Front Endocrinol (Lausanne), 2015. 6: p. 4.

7. Basyuni, S., et al., Systematic scoping review of mandibular bone tissue engineering. Br J Oral Maxillofac Surg, 2020. 58(6): p. 632–642.

8. Marshall, K.M., et al., Bioactive coatings on 3D printed scaffolds for bone regeneration: Use of Laponite™ to deliver BMP-2 for bone tissue engineering – progression through in vitro, chorioallantoic membrane assay and murine subcutaneous model validation. bioRxiv, 2023: p. 1–36.

9. Marshall, K.M., et al., Bioactive coatings on 3D printed scaffolds for bone regeneration: Translation from in vitro to in vivo models and the impact of material properties and growth factor concentration. bioRxiv, 2023.

10. Girotti, A., et al., Elastin-like recombinamers: biosynthetic strategies and biotechnological applications. Biotechnol J, 2011. 6(10): p. 1174–86.

11. Elsharkawy, S., et al., Protein disorder-order interplay to guide the growth of hierarchical mineralized structures. Nat Commun, 2018. 9(1): p. 2145.

12. Hasan, A., et al., Mineralizing Coating on 3D Printed Scaffolds for the Promotion of Osseointegration. Front Bioeng Biotechnol, 2022. 10: p. 836386.

13. Cheng, Z.A., et al., Nanoscale Coatings for Ultralow Dose BMP-2-Driven Regeneration of Critical-Sized Bone Defects. Adv Sci, 2019. 6(2): p. 1800361.

14. Page, D.J., et al., Injectable nanoclay gels for angiogenesis. Acta Biomater, 2019. 100: p. 378–387.

15. Dawson, J.I., et al., Clay gels for the delivery of regenerative microenvironments. Adv Mater, 2011. 23(29): p. 3304–8.

16. Gibbs, D.M., et al., Bone induction at physiological doses of BMP through localization by clay nanoparticle gels. Biomaterials, 2016. 99: p. 16–23.

17. Xue, R., et al., Elastin-like recombinamer-mediated hierarchical mineralization coatings on Zr-16Nb-xTi (x = 4,16 wt%) alloy surfaces improve biocompatibility. Biomater Adv, 2023. 151: p. 213471.

18. Gunderson, Z.J., et al., A comprehensive review of mouse diaphyseal femur fracture models. Injury, 2020. 51(7): p. 1439–1447.

19. Rocha, T., et al., PTH1-34 improves devitalized allogenic bone graft healing in a murine femoral critical size defect. Injury, 2021. 52 **Suppl 3**: p. S3–S12.

20. Findeisen, L., et al., Cell spheroids are as effective as single cells suspensions in the treatment of critical-sized bone defects. BMC Musculoskelet Disord, 2021. 22(1): p. 401.

21. Manassero, M., et al., A novel murine femoral segmental critical-sized defect model stabilized by plate osteosynthesis for bone tissue engineering purposes. Tissue Eng Part C Methods, 2013. 19(4): p. 271–80.

22. Liu, K., et al., A murine femoral segmental defect model for bone tissue engineering using a novel rigid internal fixation system. J Surg Res, 2013. 183(2): p. 493–502.

23. Tiyapatanaputi, P., et al., A novel murine segmental femoral graft model. Journal of Orthopaedic Research, 2004. 22: p. 1254–1260.

24. Lang, A., et al., Osteotomy models - the current status on pain scoring and management in small rodents. Lab Anim, 2016. 50(6): p. 433–441.

25. Zwingenberger, S., et al., Establishment of a femoral critical-size bone defect model in immunodeficient mice. J Surg Res, 2013. 181(1): p. e7–e14.

26. Uusitalo, H., et al., A Metaphyseal Defect Model of the Femur for Studies of Murine Bone Healing. Bone, 2001. 28(4): p. 423–429.

27. Kanczler, J.M., et al., The effect of the delivery of vascular endothelial growth factor and bone morphogenic protein-2 to osteoprogenitor cell populations on bone formation. Biomaterials, 2010. 31(6): p. 1242–50.

28. Chung, I., et al., Syntheses and evaluation of biodegradable multifunctional polymer networks. European Polymer Journal, 2003. 39(9): p. 1817–1822.

29. Green, B.J., et al., Effect of Molecular Weight and Functionality on Acrylated Poly(caprolactone) for Stereolithography and Biomedical Applications. Biomacromolecules, 2018. 19(9): p. 3682–3692.

30. Field, J., et al., A Tuneable, Photocurable, Poly(Caprolactone)-Based Resin for Tissue Engineering-Synthesis, Characterisation and Use in Stereolithography. Molecules, 2021. 26(5): p. 1199.

31. Bobbert, F.S.L., et al., Additively manufactured metallic porous biomaterials based on minimal surfaces: A unique combination of topological, mechanical, and mass transport properties. Acta Biomater, 2017. 53: p. 572–584.

32. Alba-Perez, A., et al., Plasma polymerised nanoscale coatings of controlled thickness for efficient solid-phase presentation of growth factors. Materials Science and Engineering: C, 2020. 113: p. 110966.

33. Clough, B.H., M.R. McCarley, and C.A. Gregory, A simple critical-sized femoral defect model in mice. J Vis Exp, 2015. e52368(97).

34. Marshall, W.G., et al., Bioengineering an Osteoinductive Treatment for Bone Healing Disorders: A Small Animal Case Series. VCOT Open, 2023. 06(01): p. e41–e51.

35. Yu, Y.Y., et al., Bone morphogenetic protein 2 stimulates endochondral ossification by regulating periosteal cell fate during bone repair. Bone, 2010. 47(1): p. 65–73.

36. Kanczler, J.M., et al., Biocompatibility and osteogenic potential of human fetal femur-derived cells on surface selective laser sintered scaffolds. Acta Biomater, 2009. 5(6): p. 2063–71.

37. Kanczler, J.M., et al., The effect of mesenchymal populations and vascular endothelial growth factor delivered from biodegradable polymer scaffolds on bone formation. Biomaterials, 2008. 29(12): p. 1892–900.

38. Wehrle, E., et al., Evaluation of longitudinal time-lapsed in vivo micro-CT for monitoring fracture healing in mouse femur defect models. Sci Rep, 2019. 9(1): p. 17445.

39. Kaur, A., S. Mohan, and C.H. Rundle, A segmental defect adaptation of the mouse closed femur fracture model for the analysis of severely impaired bone healing. Animal Model Exp Med, 2020. 3(2): p. 130–139.

40. Augat, P., M. Hollensteiner, and C. von Ruden, The role of mechanical stimulation in the enhancement of bone healing. Injury, 2021. 52 **Suppl 2**: p. S78–S83.

41. Hente, R.W. and S.M. Perren, Tissue deformation controlling fracture healing. J Biomech, 2021. 125: p. 110576.

42. Windolf, M., et al., The relation between fracture activity and bone healing with special reference to the early healing phase - A preclinical study. Injury, 2021. 52(1): p. 71–77.

43. Schreivogel, S., et al., Load-induced osteogenic differentiation of mesenchymal stromal cells is caused by mechano-regulated autocrine signaling. Journal of Tissue Engineering and Regenerative Medicine, 2019. 13(11): p. 1992–2008.

44. Claes, L., Improvement of clinical fracture healing - What can be learned from mechano-biological research? J Biomech, 2021. 115: p. 110148.

