## Supplemental information for "Bioactive coatings on 3D printed polycaprolactone scaffolds for bone regeneration: a novel murine femur defect model for examination of the biomaterial capacity for repair"

### Supplementary information

#### Intramedullary pin preparation

To create the IM pin, a sterile 25-gauge needle was threaded through the EO sterile scaffold and cut with pliers (sterilised in 70% ethanol) to 1.2 cm in length. In the second pilot study, polyimide/plastic tubing, was used to aid µCT imaging of the femur and subsequent data analysis, with n=4 per group of metal pin versus plastic pin. The metal needle/pin was used as a comparator and served as an additional control in case there were adverse effects from the PCL-TMA900 scaffold rather than from the use of a polyimide pin (**Supplementary Figure 1**).

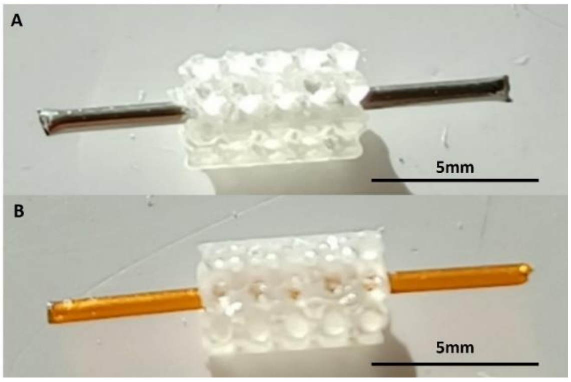


**S. Figure 1:** The metal and polyimide/plastic pin. (A) The metal pin within the PCL-TMA900 scaffold and (B) the polyimide/plastic pin within the PCL-TMA900 scaffold. Length of pins 1.2 cm and PCL-TMA900 scaffolds 5 mm, scale bar 5 mm.

#### First pilot study results – metal IM pin application, validation of defect length and confirmation of bone formation in response to collagen sponge/BMP-2 as a positive control construct

In the first pilot study, extrusion printed 5 mm long cylindrical or cuboidal PCL scaffolds were used due to known material biocompatibility and secured with a metal pin. All mice recovered uneventfully, with full normal range of movement immediately after surgery of the operated left limb. The µCT scans from the day following surgery showed adequate pin placement with the pin within the medulla and there was stability of the fracture site. The collagen sponge was mineralised from the week 2 µCT scan with increased mineralisation seen at weeks 4, 6 and 8, confirming suitability as a positive control, with minimal bone formation on or within the PCL scaffolds (**Supplementary Figure 2**).

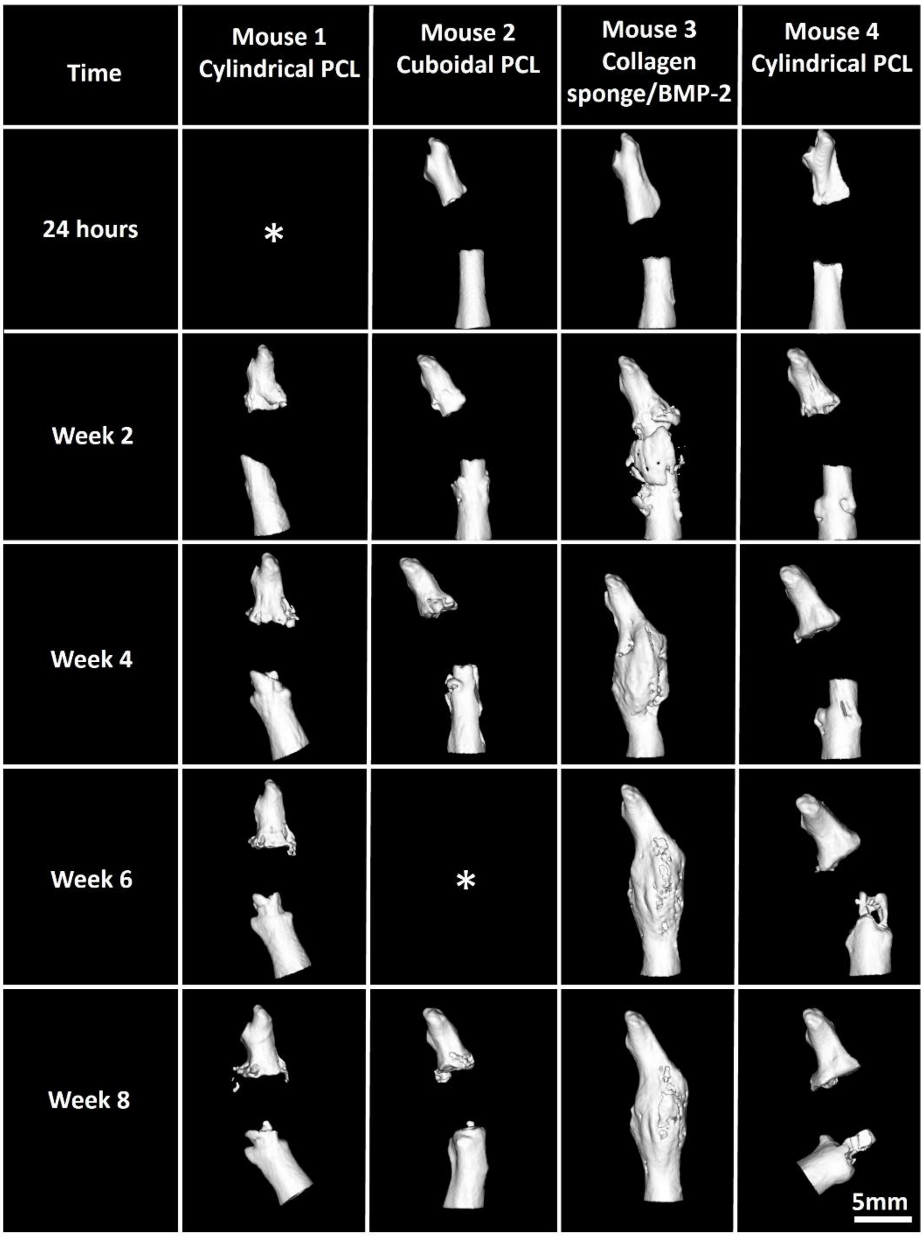


**S. Figure 2:** 3D µCT reconstructions of each limb (shown from the cranial aspect of the limb) illustrate the bone formation over time in each mouse. The periosteal reaction distal and proximal to the transected bone can be seen at week 2, with mineralisation of the collagen sponge in Mouse 3. Over time the peripheral bone from the periosteum reached the scaffolds but there was no ingrowth of bone from the marrow cavity. Mouse 4 displayed resorption of bone in the distal segment of femur and subsequent displacement of the femur. (*) Mouse 1 at 24 hours and mouse 2 at week 6 displayed movement artefacts and therefore could not be analysed at these time points. Scale bar 5 mm.

The µCT scans performed at week 8 of *ex vivo* limbs and reconstructed in 3D demonstrated marked bone formation in mouse 3 with the collagen sponge and BMP-2, while mice 1, 2 and 4 have minimal bone formation in the defect site, with the metal needle clearly visible (**Supplementary Figure 3**).

Mouse 4 at week 8 displayed images in which the bone appeared to have lysed around the needle, with the needle at a different angulation from the other 3 mice as per images between weeks 4 and 8. Therefore, displacement of the distal femur relative to the needle had occurred, necessitating the study to end for this mouse 3 days early due to inability to advance the limb normally (**Supplementary Figure 3**).

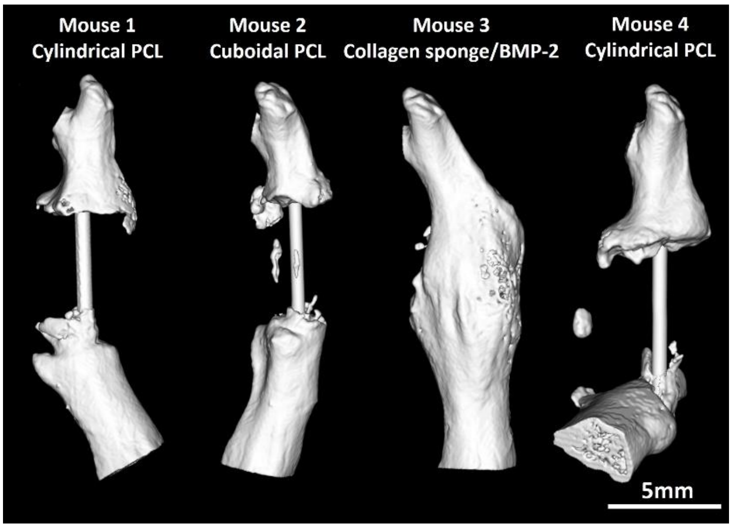


**S. Figure 3:** Week 8 *ex vivo* µCT scans of the first pilot mouse femur defect study. Mouse 1 with a cylindrical PCL scaffold, mouse 2 with a cuboidal PCL scaffold, mouse 3 with a collagen sponge with BMP-2 and mouse 4 with a cylindrical PCL scaffold. The collagen sponge with 5 µg BMP-2 resulted in bridging across the defect site and remodelling, especially at the craniomedial aspect of the bone in mouse 3, while the needle is still clearly visible in mice 1, 2 and 4. The defect did not heal in mice 1, 2 or 4 with minimal mineralisation within the PCL scaffold. Mouse 4 had displacement of the distal femur compared to the other 3 mice, that demonstrated no complications over the 8-week study period. Scale bar 5 mm.

### Second pilot study analysis: metal and plastic IM pin application and validation of PCL-TMA900 scaffold biocompatibility

In the second pilot study, there was marked bone formation observed around or within the PCL-TMA900 scaffolds on µCT imaging at week 8 (**Supplementary Figure 4 A**). The collagen sponge/BMP-2 construct in the two mice was found to be mineralised, confirming repeatability. Upon observation of the µCT data, the limbs with the plastic pin displayed a smoother, greater bone volume formed than the limbs containing a metal pin, however, when the percentage increase in bone volume for the 3 mice with the PCL-TMA900 scaffold from each group (i.e. excluding the mouse with the collagen sponge to limit variability) were analysed, the difference between the use of a metal or plastic IM pin was found to not reach significance (**Supplementary Figure 4 B**).

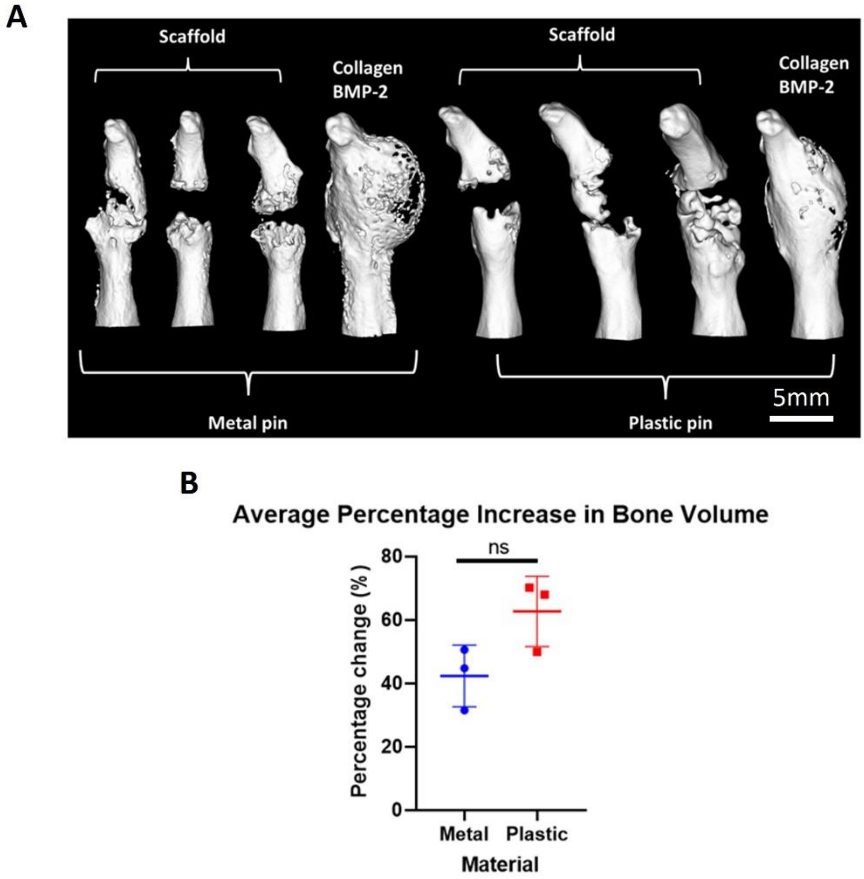


**S. Figure 4:** µCT images of bone formation at week 8 and the difference in bone formation using a metal vs plastic pin. (A) The metal pin appeared to give irregular, rough bone formation compared to the plastic pin despite the use of the copper filter and gauss filter during image acquisition and analysis respectively. N=8, scale bar 5 mm. (B) The difference in bone volume formed between metal and plastic pin fixation. Two-tailed unpaired t-test with Welch's correction performed, ns; not significant, mean and S.D. shown, n=3.

### Design of the 3D printed PCL-TMA900 scaffold

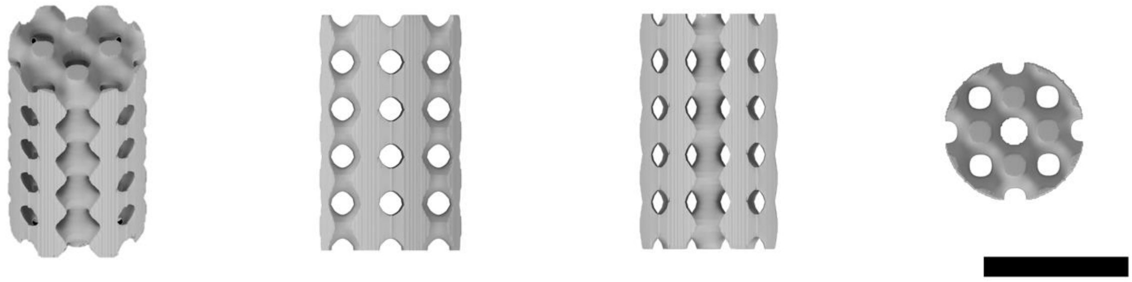

**S. Figure 5:** CAD model showing various views of the 3D printed scaffold. Height = 5 mm, diameter = 3 mm, central hole = 0.6 mm, pore size = 0.5 mm. Scale bar = 3 mm.

### Method of coating PCL-TMA900 scaffolds, clearing the marrow cavity and surgical procedure

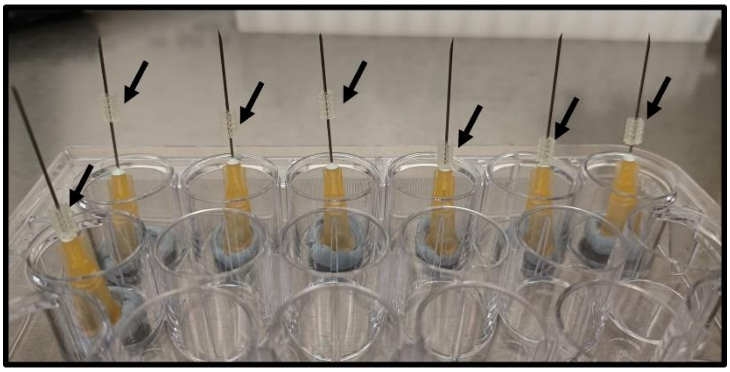

**S. Figure 6:** The PCL-TMA900 scaffolds were dried within a sterile category 2 hood. The scaffolds (black arrows) were held through the central hole by 25-gauge needles held vertically by Blu Tack in a 24 well plate, to allow the scaffolds to dry without touching a surface around the periphery.

### BMP-2 acidic buffer solution constituents

The buffer solution for dilution of Medtronic InductOs® BMP-2 for use on the collagen sponge was made by dissolving the following listed in **Table 1** in deionised water:

**Table 1:** InductOs® BMP-2 buffer solution constituents.

| **Reagent** | **Concentration** | **Quantity for 500 mL** |
|---|---|---|
| Sucrose | 0.5% | 2500 mg |
| Glycine | 2.5% | 12500 mg |
| L-Glutamic acid | 0.37% | 1850 mg |
| Sodium chloride | 0.01% | 50 mg |
| Polysorbate 80 | 0.01% | 50 mg |

### Surgical procedure for creating the femur defect

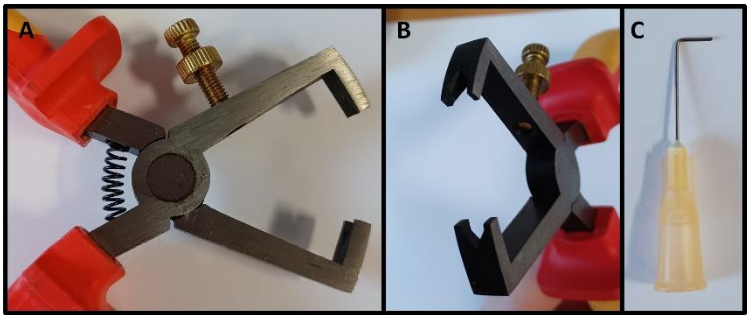


**S. Figure 7:** Preparation of 25-gauge needles for clearing of the marrow cavity. (A) and (B) wire stripping pliers were used to cut the end of a 25-gauge needle to allow the needle to be cut without crushing the end of the needle, therefore keeping the needle smooth and hollow. (C) The needle was bent at 4 mm from the cut end to allow standardised clearing of 4 mm of marrow cavity from the proximal and distal ends of the transected femur.

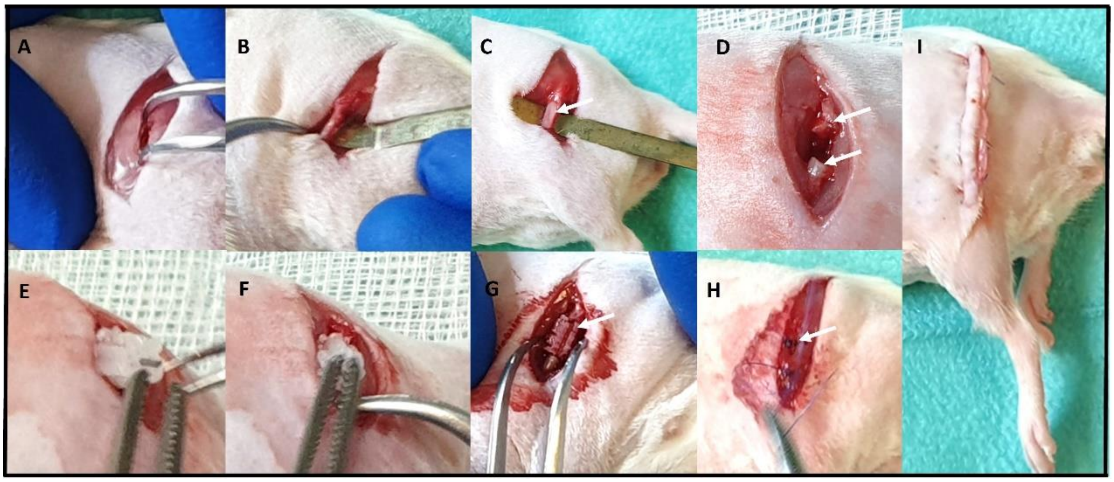


**S. Figure 8:** Mouse femur defect surgical procedure. (A) Incision of the fat/fascia overlying the femur and blunt dissection of the surrounding muscle. (B) Removal of soft tissue from around the femur while making a space for the micro-spatula underneath. (C) The micro spatula placed underneath the femur (arrow) to protect the surrounding soft tissues. (D) The two cut ends of the femur (arrows) with a gap between them. (E) Insertion of the cylindrical PCL scaffold on a 25 gauge 1.2cm needle into the proximal end of the femur. (F) Manipulation of the distal end of the femur to place the needle within the medulla of the distal section of the bone. (G) The scaffold (arrow) *in situ* with femur diaphysis above and below. (H) Closure of the muscle layer to reduce dead-space with simple interrupted sutures (arrow). (I) Closure of the skin with the everted skin edge creating a barrier from interference with the sutures by the mouse.

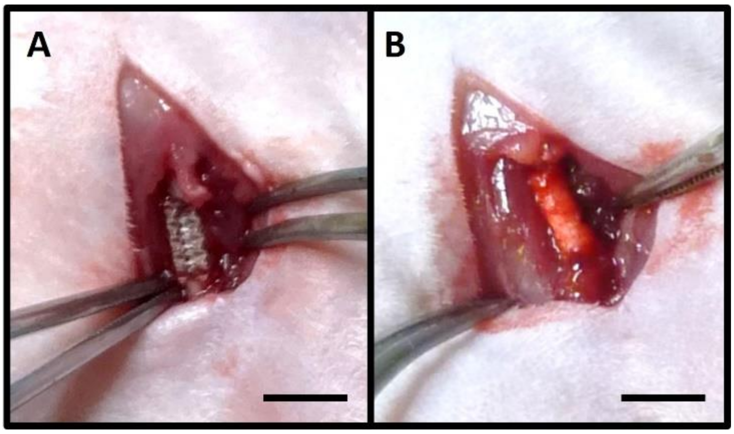


**S. Figure 9:** The materials in the femur defect site. (A) The PCL-TMA900 scaffold and, (B) the collagen sponge with 5µg of BMP-2 in position during surgery. Scale bar 5 mm.

### µCT settings for the MILabs OI-CTUHXR preclinical imaging scanner

**Table 2:** Settings used for µCT scanning of mice and *ex vivo* limbs in the femur defect studies

| **Imaging reference** | **Samples *in vivo* in rat bed** | **Cu filter settings due to metal pin (pilot studies)** | ***Ex vivo* samples in mouse bed** |
|---|---|---|---|
| Voltage (kVp) | 55 | 65 | 50 |
| Current (mA) | 0.17 | 0.13 | 0.21 |
| Exposure time (ms) | 75 | 100 | 75 |
| Filter (µm thickness) | Al 400+100 | Al100+Cu100 or Cu200 | Al 400+100 |
| Scan angle (°) | 360 | 360 | 360 |
| Step angle (°) | 0.25 | 0.25 | 0.25 |
| Pixel size | 40 | 40 (20 in *ex vivo* samples) | 20 |
| Reconstruction voxel size ($\mu m^3$) | 40 | 40 (20 in *ex vivo* samples) | 20 |
| Frame averaging | 1 | 1 | 1 |

### Bone volume analysis

A gauss filter of 2.0 was applied and the phantom was analysed in each scan. The average value of the lower density phantom (1.5 cm × 4.5 cm cylindrical area) was used to set the threshold for analysis. Initially in all studies, the bone volume to be quantified was segmented out by thresholding for the density of bone and then allocating areas of the skeleton to different groups/classes e.g. 'skeleton', 'limb', 'to analyse' and the 3D drawing tool in the software was used to exclude unwanted areas in the 2D or 3D views e.g., the pin within the collagen sponge could only be accessed on the 2D views and allocated to the volume to be discarded by this method. The region to be quantified was initially from the proximal extent around the neck of the femur to distally above the trochlear groove of the femur. The metal pin, scatter artefact and areas not to be quantified were removed from the bone volume quantification. In the case of the plastic pin, no metal pin was present to create an artefact, therefore quantification was more accurate (**Supplementary Figure 10)**.

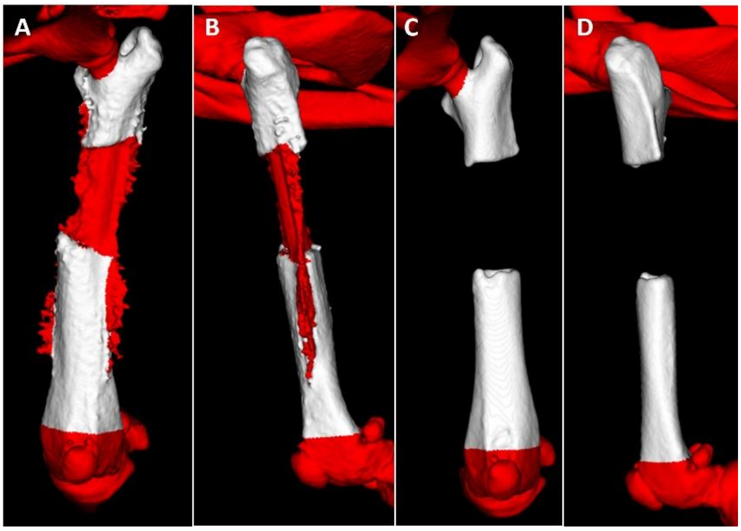


**S. Figure 10:** Segmentation of the area of interest in study 2. (A) The cranial view with the areas of interest in white and discarded areas in red, with artefact from the metal pin seen as uneven and jagged red areas. (B) Lateral view of the limb with the cut-off at the distal femur shown, with jagged red areas discarded because of interference from the metal pin. (C) Cranial view of the plastic pin to illustrate the lack of interference due to lack of metal and improved accuracy of analysis. (D) The lateral view showing the improvement in image analysis capability using a plastic pin.

For analysis of the study involving the coated PCL-TMA900 scaffolds, the segmentation method described above was used initially at all timepoints of day of surgery, week 2, 4, 6, 8, and when the samples were scanned *ex vivo*. A gauss filter of 1.0 was used as there was no metal to cause interference.

Analysis was repeated by selecting a 4 mm by 6 mm cylindrical area over the region of the scaffold (scaffold size 3 mm × 5 mm) which therefore included a 0.5 mm periphery around scaffold. The centre of the scaffold was measured to be 2.5 mm from the femur ends and 1.5 mm from the perceived edge of the scaffold. This was easiest in *ex vivo* images due to enhanced detail of the radiolucent shape of the scaffold against the surrounding soft tissues in the images (**Supplementary Figure 11**).

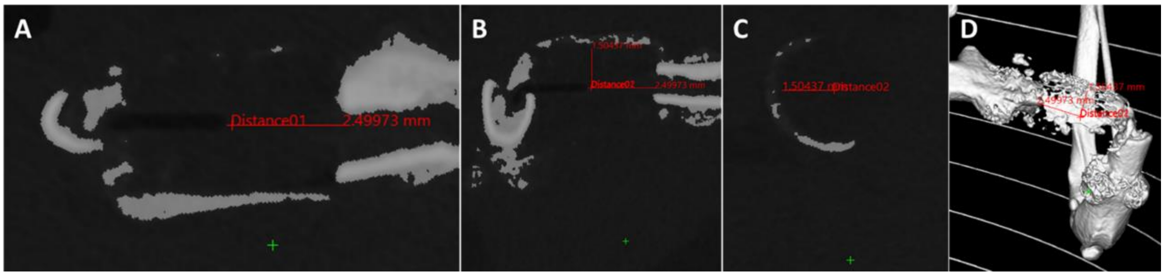


**S. Figure 11:** Cylinder placement for bone quantification. (A) The sagittal plane showing the distance to the centre point of the 5 mm scaffold. (B) The coronal plane showing 1.5 mm from the centre of the plastic pin to the outer edge of the scaffold. (C) Transverse view of the scaffold with the 1.5 mm radius marked. (D) The 3D view illustrating the 3 measurements. For the collagen sponge, the 4 mm x 6 mm cylinder was used however 0.5 mm of either end of the femur was included in analysis as this was equivalent to the limbs with the scaffold material to avoid underestimation of bone volume.

For the 4 mm × 6 mm analysis method, the scaffold was aligned in the 3 planes in 2D images and measurements were taken to determine the centre of the scaffold and the cylindrical 4 mm × 6 mm volume selected (**Supplementary Figure 12 A**). However, the bone volume would be difficult to equivocate to the scaffold as the diameter of the sponge swelled to greater than 4 mm and so the outer bone was disregarded (**Supplementary Figure 12 B**).

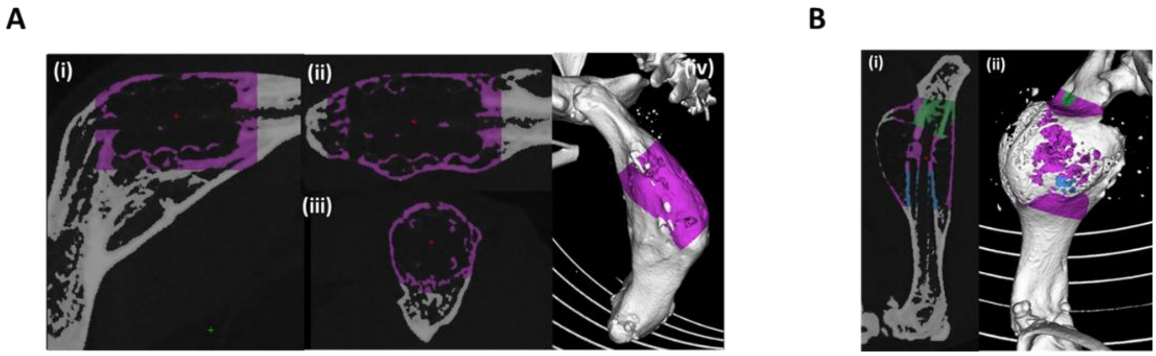


**S. Figure 12:** Collagen sponge bone quantification in a 4 mm × 6 mm cylinder. (A) Scaffold quantification in 4 mm × 6 mm cylinder. (i) The coronal plane, (ii) the sagittal plane, (iii) the transverse plane and, (iv) 3D image of the area of the cylinder selecting the 4 mm × 6 mm region of bone. The discrete red cross in each image is the centre of the scaffold/cylindrical area. (B) (i) The 4 mm × 6 mm cylindrical volume (purple) was quantified and the proximal (green) and distal (blue) femur regions were not included in the analysis of bone volume. (ii) The image in (i) shown in 3D illustrating the areas of bone wider than 4 mm diameter from the plastic pin which were excluded from analysis as they were outside the 6 mm diameter cylindrical area.

To analyse the volume of the scaffold itself a 3 mm × 5 mm cylinder was selected. The 3 mm × 5 mm cylinder was applied and quantified in the *ex vivo* images, as there was no bone present at the day of surgery within this area, and thus the volume of bone formation could be calculated without the confounding factor of the medial arch of bone which formed in the mice with scaffolds from week 2 onwards. A 3 mm × 5 mm cylinder was placed within the 6 mm × 4 mm cylindrical volume for quantification (**Supplementary Figure 13 A**). For the collagen sponge, a 3 mm × 5 mm cylindrical volume was selected, and the femur ends removed from the quantification of bone as the sponge was usually compressed to bring the femur within the 3 mm × 5 mm volume, which would lead to overestimation of bone formation. Again, the outer bone which formed wider than 3mm diameter was disregarded (**Supplementary Figure 13 B**).

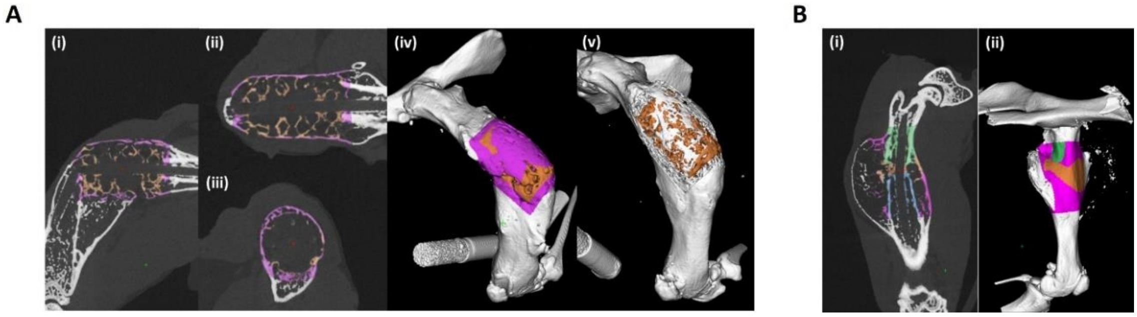


**S. Figure 13:** Scaffold and collagen sponge bone quantification using the 3 mm × 5 mm cylinder method. (A) Scaffold bone volume quantification; (i) Coronal view, (ii) sagittal view, (iii) transverse view and, (iv) 3D view of the 3 mm × 5 mm cylinder in orange within the pink 6 mm × 4 mm cylinder volume. (v) The pink 6 mm × 4 mm cylinder class has been hidden to show the quantified orange 3 mm × 5 mm cylinder of bone volume within/around the scaffold itself. (B) Collagen sponge bone quantification of a 3 mm × 5 mm cylinder volume. (i) Coronal plane view of the classes used for quantification with orange being the 3 mm × 5 mm cylinder and pink showing the previous 6 mm × 4 mm cylinder applied and the green and blue areas are the femur itself which were removed from the quantity of bone analysed. (ii) The classes seen in 3D with underestimation of volume of bone formed as the white bone protruding from the pink area was not included in quantification as it was outside the 5 mm diameter cylinder volume.

The medial and lateral sides of the most efficacious coated scaffolds; ELP/PEA/FN/BMP-2, PEA/FN/BMP-2 and Laponite/BMP-2 coated scaffolds were quantified through application of a cylinder medial and lateral within the 3 mm × 5 mm scaffold cylinder volume as the program could only produce certain standardised shapes and to ensure methodological consistency. The cylinder was positioned over the medial and lateral scaffold respectively by accurately lining up the image in the three 2D planes. The medial and lateral volumes could be segmented out and quantified individually and allowed greater visualisation of the shape of bone formed within the scaffold itself (**Supplementary Figure 14**).

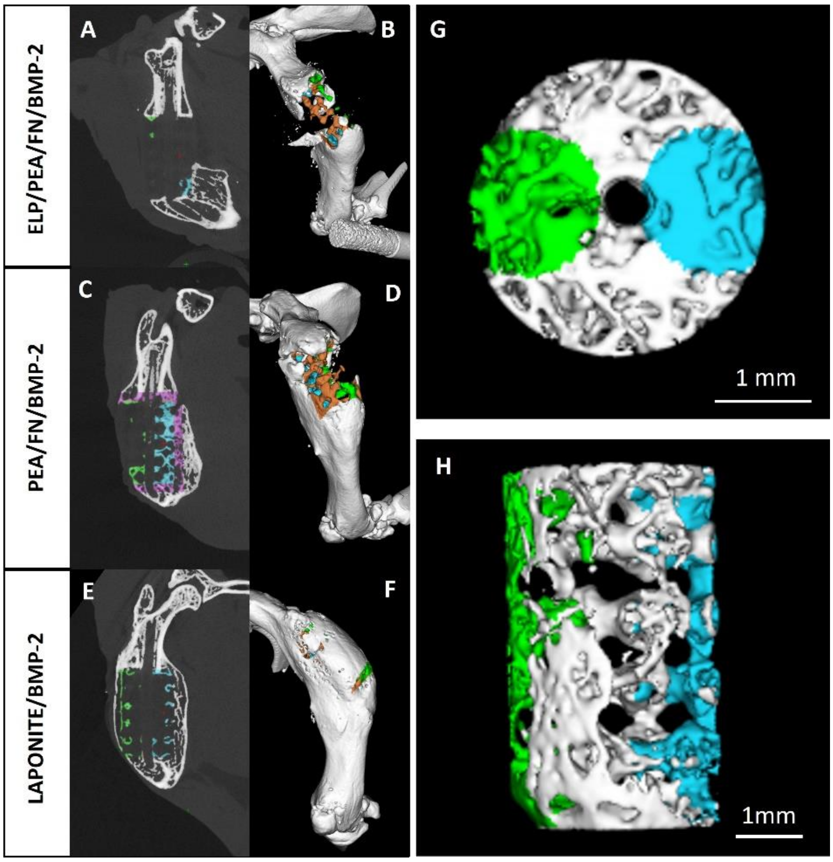


**S. Figure 14:** Quantification of the medial and lateral bone volumes. (A and B) ELP/PEA/FN/BMP-2 coated scaffold, (C and D) PEA/FN/BMP-2 coated scaffold and (E and F) Laponite/BMP-2 coated scaffold with the lateral (green) and medial (blue) bone selected within the 3 mm × 5 mm scaffold area (orange) in each image in 2D (A, C and E) and 3D (B, D and F) respectively. (G) End-on 3D view of the bone formed in the 3 mm × 5 mm volume of the Laponite/BMP-2 coated scaffold with the lateral (green) and medial (blue) bone selected, scale bar 1 mm. (H) Side-on 3D view of the same bone volume showing the bone the laterally (green) and medially (blue) and interlinked bone within the pores of the scaffold, scale bar 1 mm.

### Histological evaluation of the plastic pin

In a mouse with the PCL-TMA900 scaffold implanted, the plastic pin was observed within the section as the plastic pin could be cut with the microtome, without destroying the surrounding tissues. The new active bone was seen distal to the scaffold with fibrous and ossifying tissue around the pores of the scaffold (**Supplementary Figure 15**).

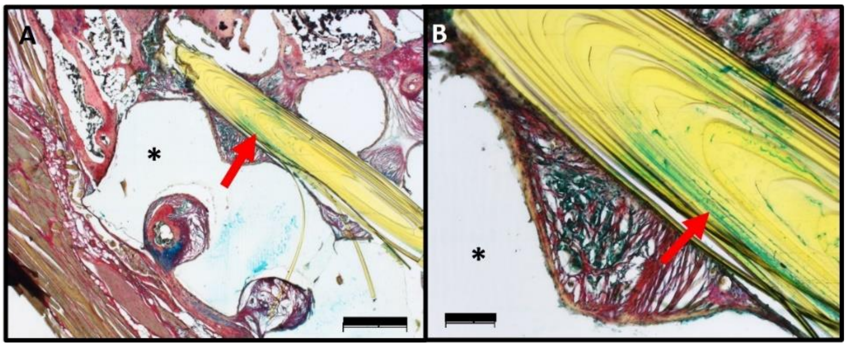


**S. Figure 15:** Histology of a PCL-TMA900 scaffold with plastic pin. (A) Alcian blue and Sirius red staining showing the plastic pin (red arrow) sectioned within the centre of the scaffold (*) scale bar 500 µm. (B) Image A at higher magnification, with the plastic pin shown (red arrow), scale bar 100 µm.

### Gross dissection images

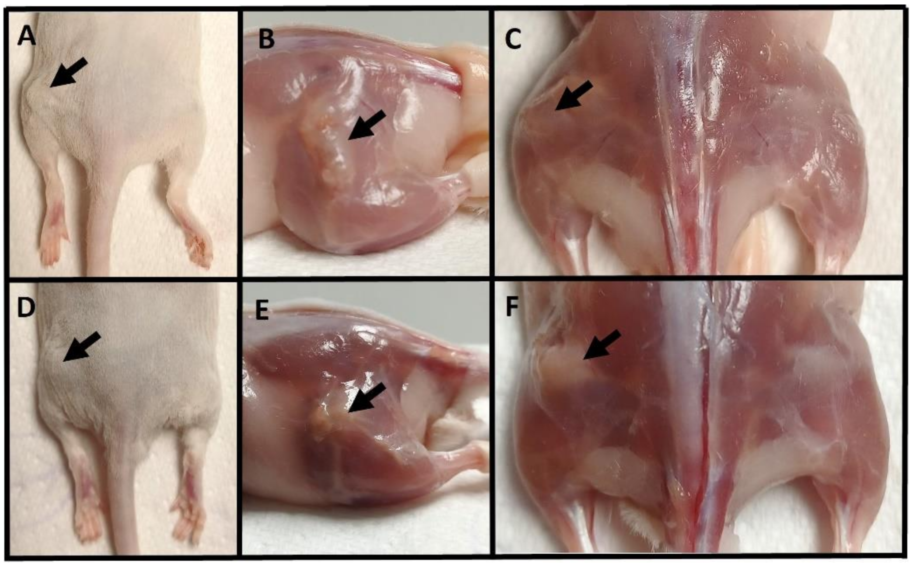


**S. Figure 16:** Gross dissection images of representative mice. (A-C) PEA/FN/BMP-2 coated PCL-TMA900 scaffold (A) the scaffold edge could be felt under the skin (arrow), (B) the scaffold (black arrow) can be seen sitting at ~90° to the ground rather than at the normal ~45° but is firmly between the ends of the transected femur (C) The scaffold (arrow) shown from above. (D-F) Collagen/BMP-2 construct (D) the callus surrounding the collagen sponge could be palpated as a firm, round mass under the skin (arrow), (E) the white callus of bone surrounding the collagen sponge (black arrow) can be seen, (F) the white callus around the collagen sponge (arrow) shown from above.

Analysis of the coated PCL-TMA900 scaffolds and collagen

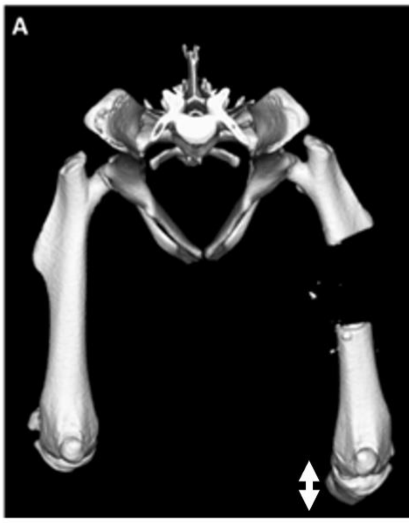


**S. Figure 17:** Visualisation by µCT of the lengthening of the limb. The limb orientation was correct on the day of surgery, however the limb was longer on the operated side (double-headed arrow) prior to bending of the pin.

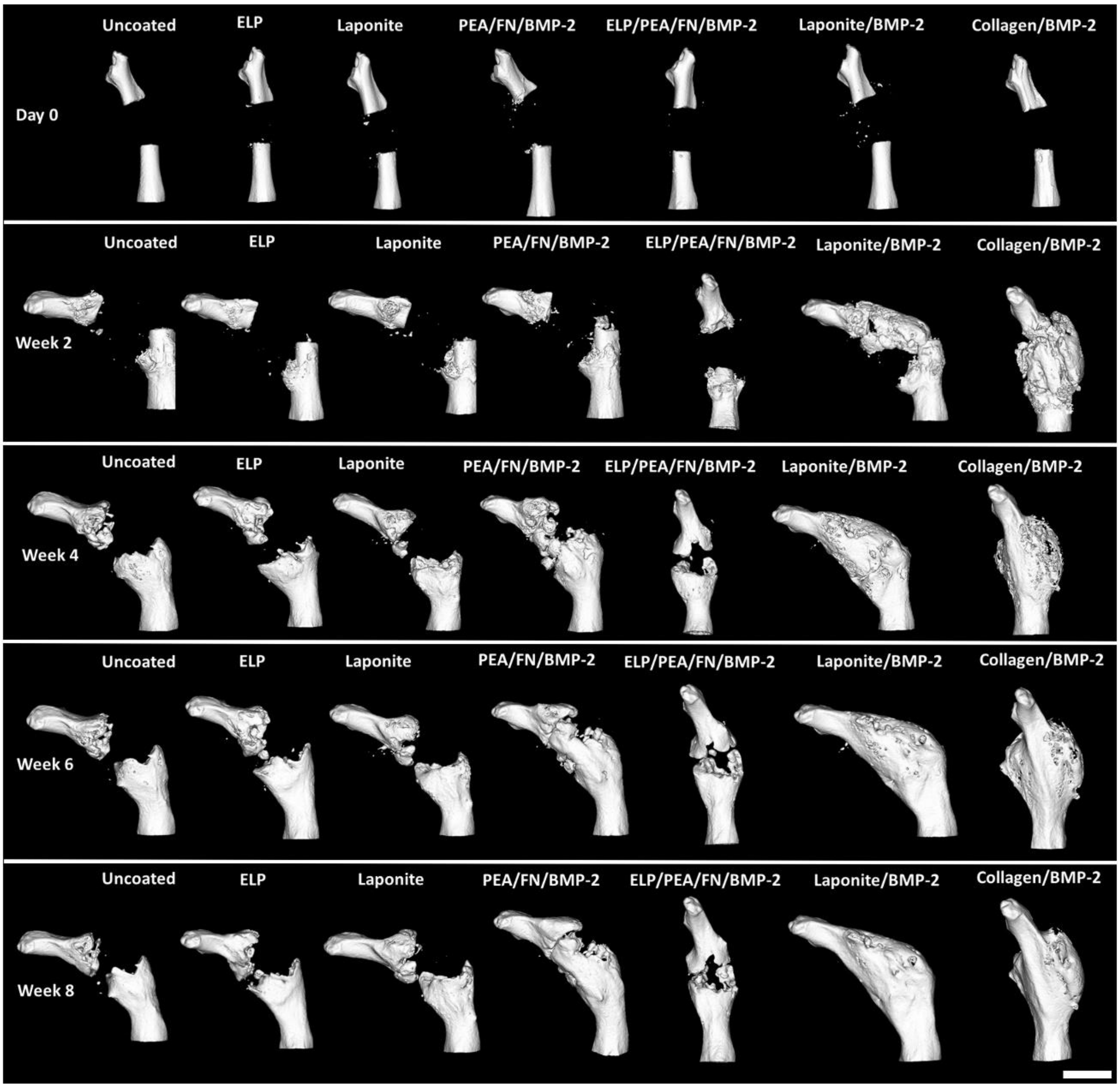

**S. Figure 18:** Cranial view of the femurs using µCT showing changes in bone volume observed over time. From the initial scans the pin bent and remained at the same angle from week 2 onwards. This resulted in an inward, medial bend to the limb in all mice except the ELP/PEA/FN/BMP-2 group. Over time the Laponite/BMP-2 coating displayed the greatest bone healing compared to the other coating groups. The collagen/BMP-2 group displayed complete union of the fracture site with smooth remodelling of the cortex craniomedially. Representative images shown from each group (n=6, n=5 for ELP/PEA/FN/BMP-2 after week 2), scale bar 5 mm.

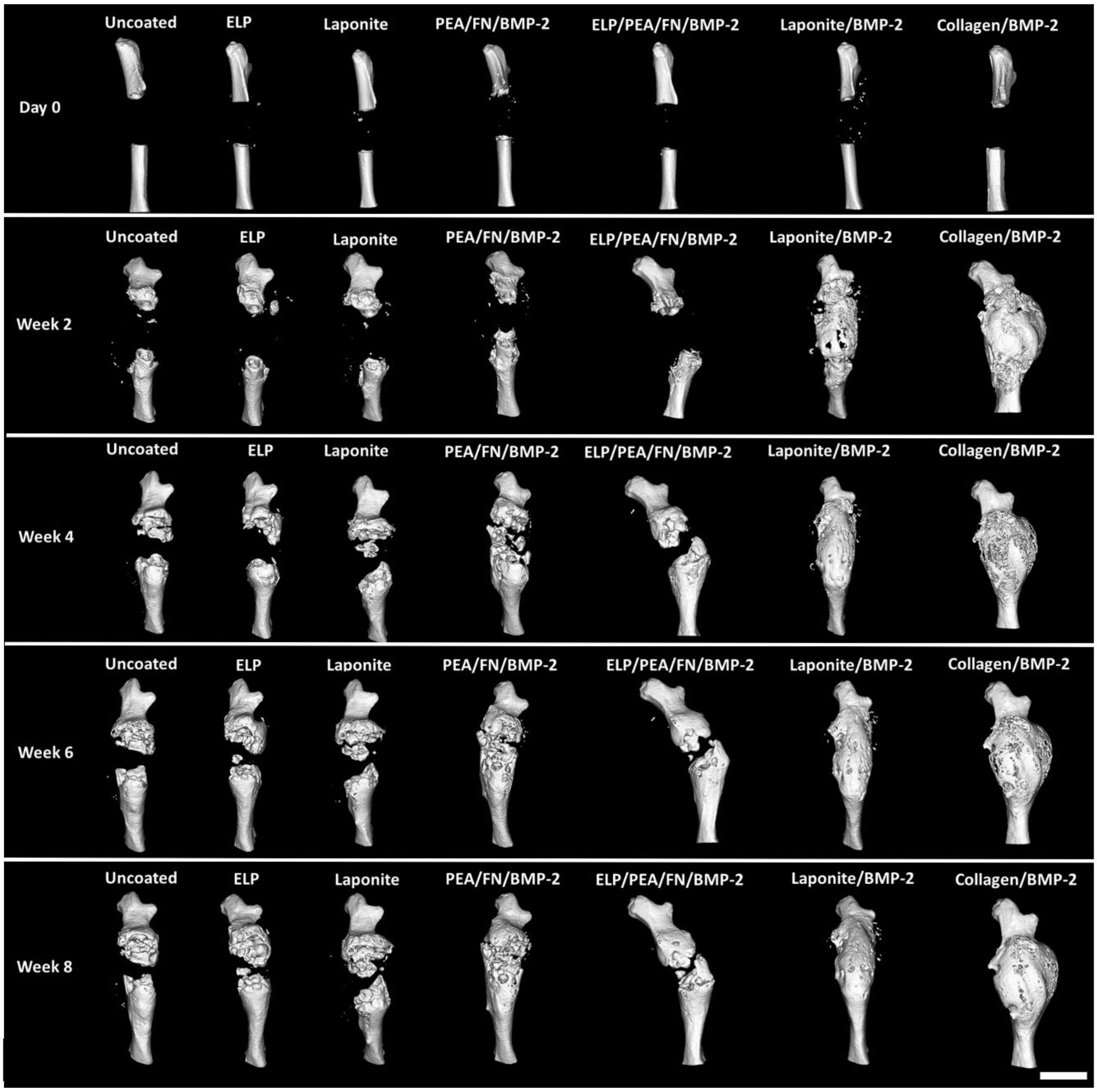


**S. Figure 19:** Lateral view of the femurs using µCT to show bone volume change over time. From the initial scans, the femurs were aligned in a cranial/caudal orientation, except the ELP/PEA/FN/BMP-2 group in which the scaffold moved caudally and as a consequence the distal limb displaced cranially to a degree. Over time the Laponite/BMP-2 coating displayed the greatest bone healing compared to the other coating groups. The collagen/BMP-2 group displayed complete union of the fracture site with a spherical shell of bone laterally and caudally. Representative images shown from each group (n=6, n=5 for ELP/PEA/FN/BMP-2 after week 2), scale bar 5 mm.

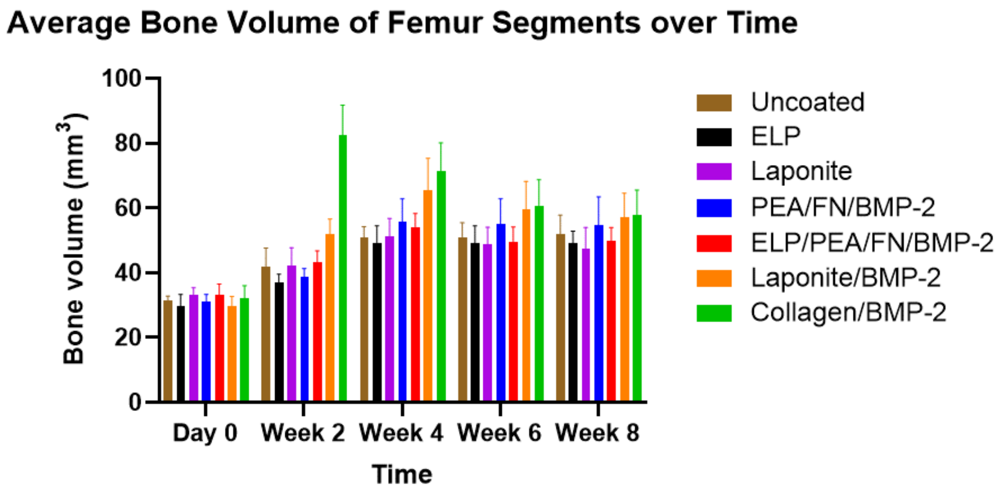


**S. Figure 20:** Average bone formation in each group. A marked increase in bone volume in the collagen sponge/BMP-2 group at week 2, followed by a reduction over the subsequent six weeks was observed. The coated scaffold bone volumes increased to week 4 and subsequently were observed to plateau. N=6 each group, except n=5 for ELP/PEA/FN/BMP-2 group, mean and S.D. shown.

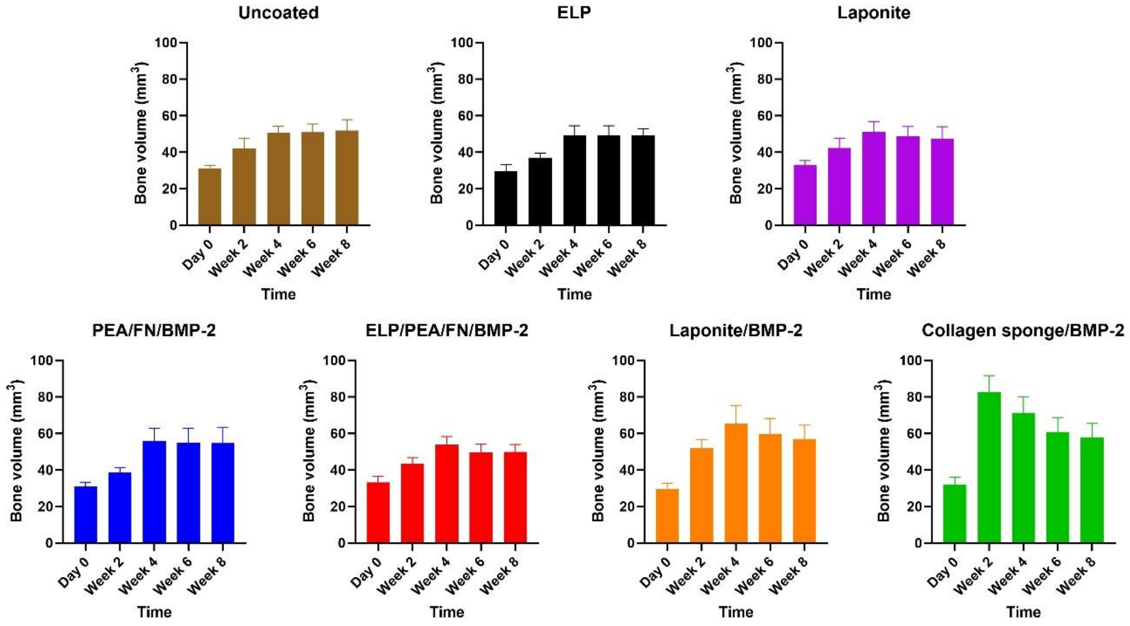


**S. Figure 21:** Average bone volume data of femur segments over time for each coating or material. Each group showed an increase in bone volume to week 4 followed by a reduction or stasis of the bone volume formed at weeks 6 and 8. The collagen sponge displayed an initial large increase in bone volume followed by reduction in volume over the remaining 6 weeks of the study. N=6 each group, except n=5 for ELP/PEA/FN/BMP-2 group, mean and S.D. shown.

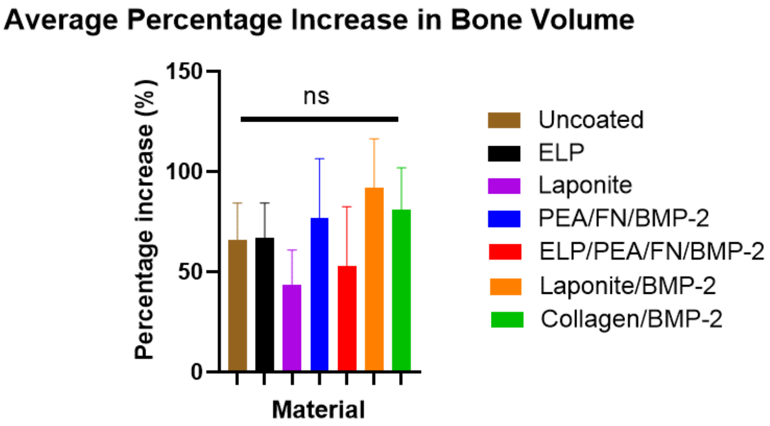


**S. Figure 22:** Percentage increase in bone volume in each segment. Averages of percentage increase for each group had large standard deviations due to the angulation of the femur n=6 all groups except n=5 for ELP/PEA/FN/BMP-2 group. N=6 each group, except n=5 for ELP/PEA/FN/BMP-2 group, one-way ANOVA with Dunnett's multiple comparison test, ns; not significant, mean and S.D. shown.

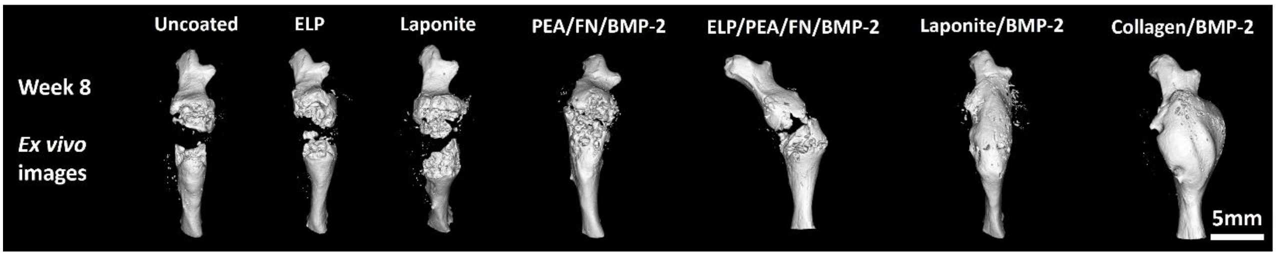


**S. Figure 23:** *Ex vivo* 3D µCT images from lateral aspect of the limbs. The Laponite/BMP-2 coating on the PCL-TMA900 scaffold showed union of the bone defect with a more streamlined shape due to the coating being on the scaffold compared to the uncontrolled swelling of the collagen sponge to form a large callus. Representative images shown from each group (n=6, n=5 for ELP/PEA/FN/BMP-2 after week 2), scale bar 5 mm.

### Overview histology of each scaffold group assessed

Slides were stained with Alcian blue and Sirius red per the methods section and were imaged using the Virtual Slide System VS110 microscope (Olympus, Japan) at 10 × magnification and images taken using VS Desktop software (Olympus, Japan) to produce an overview of the representative limb from each group (**Supplementary Figure 24**).

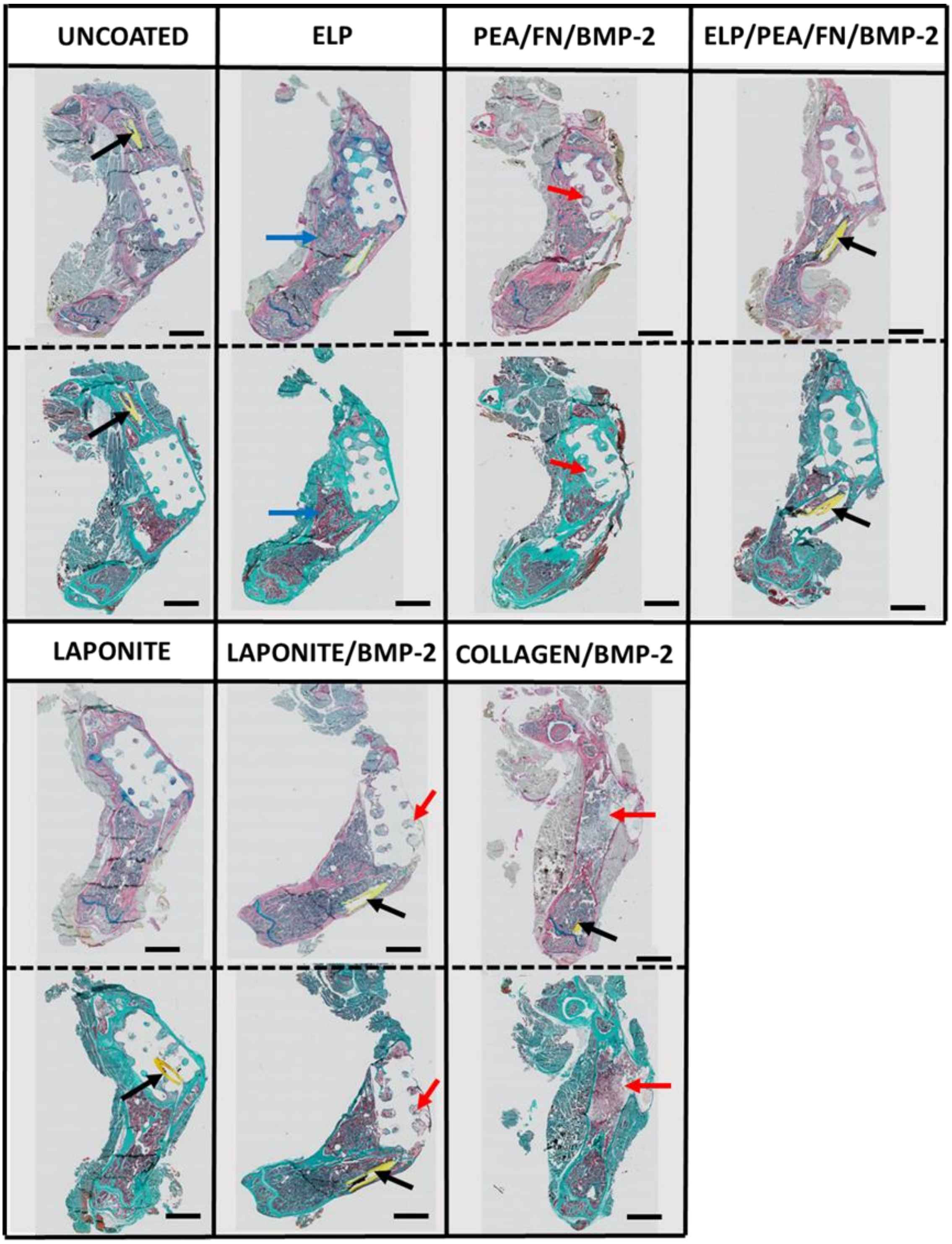


**S. Figure 24:** Histology sections of representative limbs from each scaffold group. Alcian blue/Sirius red staining of top rows and Goldner's trichrome staining of bottom rows. The curvature of the limb can be seen in each scaffold group, with the plastic pin (black arrow) seen as a yellow structure within the medullary canal. The medial shelf of new bone (blue arrow) is seen. Bone formed within the pores of the scaffold (red arrow) in PEA/FN/BMP-2 and Laponite/BMP-2 examples and within the collage sponge (red arrow) can be seen, while fibrous tissue is present surrounding the scaffold or within the scaffold pores in the other groups. Scale bar 2 mm.
